# Organisation of gene programs revealed by unsupervised analysis of diverse gene-trait associations

**DOI:** 10.1101/2022.04.07.487559

**Authors:** Dalia Mizikovsky, Marina Naval Sanchez, Christian M. Nefzger, Gabriel Cuellar Partida, Nathan J. Palpant

**Author notes:** Co-corresponding authors, **Corresponding authors**, Nathan Palpant, Institute for Molecular Bioscience, University of Queensland Brisbane, QLD, Australia, E, Gabriel Cuellar Partida Diamantina Institute University of Queensland Brisbane, QLD, Australia, E. 23andMe Inc (Gabriel Cuellar Partida).

## Abstract

Genome wide association studies provide statistical measures of gene-trait associations that reveal how genetic variation influences phenotypes. This study develops an unsupervised dimensionality reduction method called UnTANGLeD (Unsupervised Trait Analysis of Networks from Gene Level Data) which organises 16,849 genes into discrete gene programs by measuring the statistical association between genetic variants and 1,393 diverse complex traits. UnTANGLeD reveals 173 gene clusters enriched for protein-protein interactions and highly distinct biological processes governing development, signalling, disease, and homeostasis. We identify diverse gene networks with robust interactions but not associated with known biological processes. Analysis of independent disease traits shows that UnTANGLeD gene clusters are conserved across all complex traits, providing a simple and powerful framework to predict novel gene candidates and programs influencing orthogonal disease phenotypes. Collectively, this study demonstrates that gene programs co-ordinately orchestrating cell functions can be identified without reliance on prior knowledge, providing a method for use in functional annotation, hypothesis generation, machine learning and prediction algorithms, and the interpretation of diverse genomic data.

## INTRODUCTION

Generation of consortium-scale data such as ENCODE (1), the Human Cell Atlas (2) and the UKBiobank (3) coupled with the development of advanced computational methods is enabling the creation of transformative models that harness the natural diversity of biological systems. These models draw on the relationships and patterns derived from biological data to establish quantitative frameworks that can make highly accurate predictions, with implications for nearly every field of biology. For example, in the field of structural biology, patterns in the sequences and structures of proteins’ evolutionary homologs reveal how amino acids interact, enabling prediction of protein structure with atomic accuracy (4). Similarly, patterns of repressive histone methylation (H3K27me3) across hundreds of human cell types enable identification of genes governing cell decisions and functions for any cell type and organism (5).

Genome wide association studies (GWAS) characterise the genomic variation underlying complex traits and diseases, providing insights into how genes affect biological processes (6). Despite the wealth of variant-trait association information, GWAS studies predominantly focus on elucidating the genetic basis of a single trait or a group of highly related traits (6, 7). Here, we utilize patterns of genomic variation across hundreds of diverse phenotypes as the basis for an unsupervised method to parse the organisation of gene programs in cells.

We hypothesised that complex traits are underpinned by conserved gene programs that can be identified by studying associations between genetic variation and phenotypic variation. To test this, we developed UnTANGLeD (Unsupervised Trait Analysis of Networks from Gene Level Data), which identifies patterns of association between genes and hundreds of diverse phenotypes. UnTANGLeD creates a phenotypic signature to cluster genes with similar associations across many traits in an unsupervised manner into gene programs controlling cell biological processes **(Figure 1)**.

**Figure 1.**
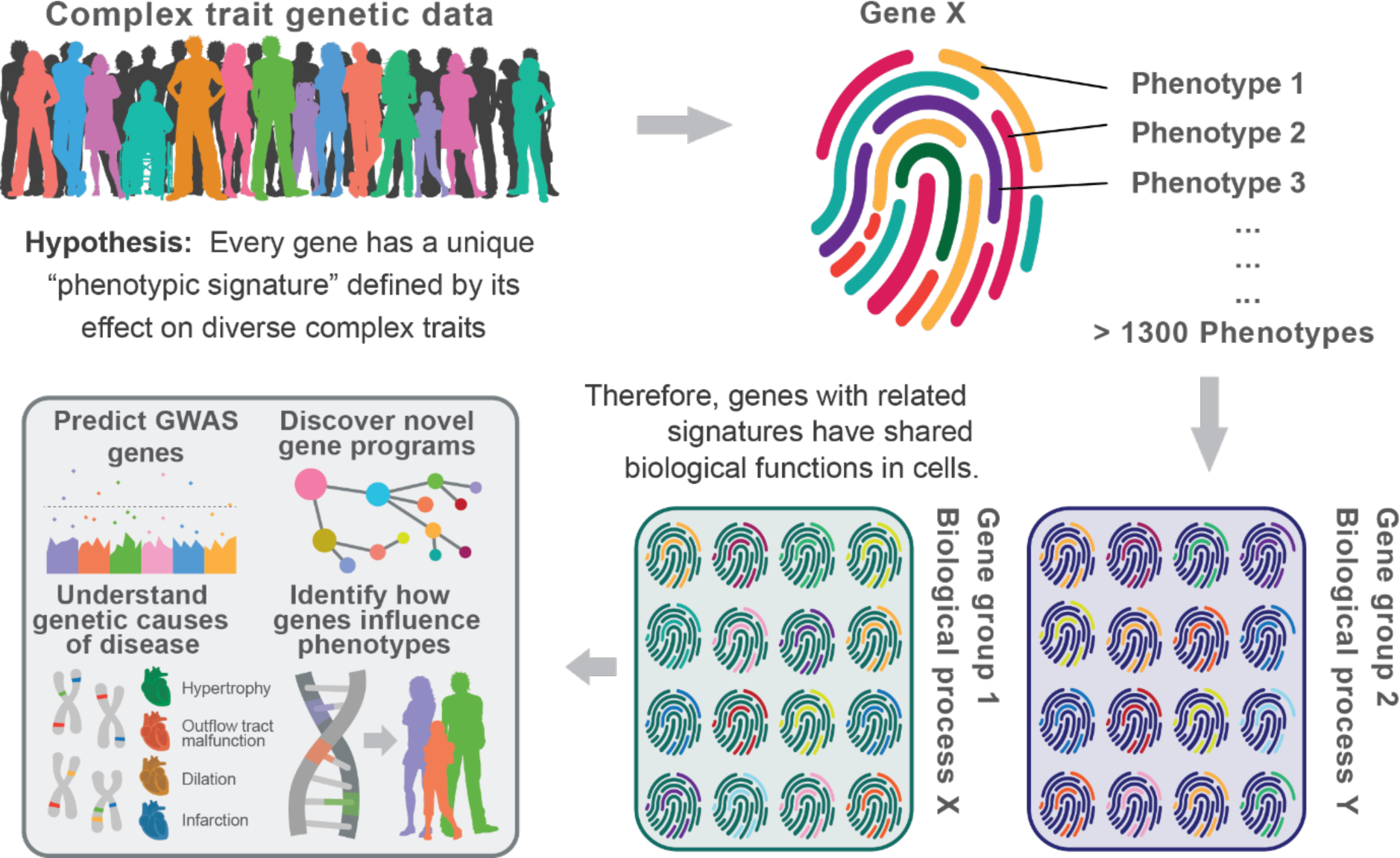
Schematic of central model design. Complex trait genetic data provide a unique association signature for each gene which can be used to parse the genome into functionally related gene sets.

We used a gene-trait association matrix derived from GWAS data for 1,393 complex traits to infer co-ordinately acting gene programs that represent both known and novel biological processes. While the scale of associations available from public GWAS data is underpowered to saturate the accuracy of our model, we demonstrate that UnTANGLeD can be applied to any orthogonal GWAS data to predict the genetic basis of disease including in underpowered and transethnic GWAS data. UnTANGLeD provides a powerful analytical framework for studies in population genetics, cell biology, and genomics, that will improve as more data emerges. Collectively, this study provides a statistical framework for defining genes orchestrating biological processes by evaluating genetic signatures across diverse complex traits.

## MATERIALS AND METHODS

### Data Collection

S-MultiXcan results for 1,393 phenotypes with statistically significant SNP-based heritability (p < 0.05) were downloaded from CTG-VL (http://vl.genoma.io). Phenotypes are listed in **Table S2**. SNP-based heritability was estimated using linkage disequilibrium score regression (LDSR). The significance values reflecting the strength of the association between each gene and trait across all tissues were compiled into a gene-trait association matrix.

### Dimensionality Reduction Analysis Pipeline

All genes with fewer than 2 significant associations across all phenotypes (p < 10^-4^) were removed, leaving 16849 genes. Following this, all values in the gene-trait association matrix were chi-squared transformed. Infinite values produced when transforming very small p-value (<1e-300) due to floating point precision were replaced with 1,415, which was 5 greater than the largest non-infinite value. The data was then normalised by the sum of chi-squared values per phenotype and scaled by a factor of 10,000. 10 principal components were retained from the principal component analysis (PCA). Clustering of genes was performed using the native *Seurat* shared-nearest neighbour algorithm. Clustering iterations were performed at increasing resolutions from 0.2 to 20 in increments of 0.2. The resolution is a parameter from Seurat where increased values lead to a greater number of clusters. Cluster assignments were compiled into a consensus distance matrix, where each gene pair had a value representing how often they were grouped together out of 100 potential matches. 100 was then subtracted from the values and they were made absolute to transform the matrix into a dissimilarity matrix. Agglomerative clustering using Ward’s minimum variance method, as implemented in the *stats* package, was applied to the consensus matrix directly. The average silhouette score (a metric used to calculate how well a data point relates to its cluster) across all genes was calculated using the *cluster* package from 2 to 300 clusters. The *inflection* package was used to calculate the plateau point, which was determined to be the optimal number of clusters. Pearson’s correlation was used to determine the correlation of a gene with the other genes in the same cluster based on chi-squared association values.

### Enrichment Analyses

GO, DO, KEGG enrichment, colocalization and tissue specificity enrichment were performed using *ClusterProfiler* (8). An FDR corrected significance value of p < 0.01 was used. Colocalization was determined using *ClusterProfiler* enrichment for the Molecular Signatures Database collection 3: positional gene sets (9). The largest proportion of genes within a cluster belonging to a single genomic region was divided by the total number of genes within the cluster to identify the maximum degree of colocalization. STRING enrichment analysis was performed using the *STRINGdb* package, with a significance threshold of p < 0.001 and a confidence threshold of 0.400. STRING enrichment analysis without the text-mining component was performed using the online STRING interface (https://string-db.org/) for clusters found to have PPI enrichment in the prior analysis with a confidence threshold of 0.150 to preserve predicted interactions reinforced by other components. For the calculation of the correlation between the loss of enrichment and the degree of colocalization, clusters 111 and 173 were removed due to having well established biological functions despite being highly colocalised. Broad enrichment analysis for more specialised gene sets was performed using *EnrichR* (https://maayanlab.cloud/Enrichr/) across all 192 libraries. Redundant libraries, including GO, KEGG, chromosomal location and NIH-grant associated libraries were excluded. The top significant term from each library for each cluster are reported in **Table S9**. A significance value threshold of 0.01, after correction for multiple testing, was used. For identification of genes possessing the same protein domains or belonging to the same family, the EnrichR library ‘Pfam_Domains_2019’ was used. A distinct protein family or domain was defined by collating the family or domain terms together that shared genes until there was no overlap between them. Protein terms did not need to be significantly enriched, but two or more members of a protein family had to be present in a single cluster.

### Permutations

Five permutations were generated by re-ordering the values within the gene-trait association matrix. These permutations were analysed as described above. A one-way ANOVA with FDR corrected pairwise comparisons was performed to identify significant differences in the number of enriched clusters, total enriched GO terms and the most significant GO enrichment of any cluster.

### Phenotype Associations

The gene-trait association matrix containing p-values was -log10 transformed. All infinite values generated due to floating point precision were windsorized with 315, which was 5 greater than the maximum finite value. The phenotypic associations for the genes within a cluster were extracted, averaged, normalised for their average associations across the dataset and ranked.

### Clustering quality in dimensionality reduction methods

We extracted the UMAP coordinates for all genes as calculated by *Seurat*. Following this, we identified the 10 closest neighbours for each gene and calculated the average correlation of chi-squared association values between the gene and its neighbours. The UMAP was re-plotted representing the average correlation with each point colour. We repeated the process, instead colouring by the number of significant associations for each gene.

### Prediction of novel genes using an underpowered GWAS of the same trait

#### Data collection and S-MultiXcan Analysis

We selected 13 phenotypes for which GWAS studies had been performed at differing cohort sizes or ethnicities for the same, or comparable traits. The specific studies and their respective details can be found in **Table S1**. Summary statistics were downloaded from various sources and harmonised using MetaXcan’s in-built harmonization (https://github.com/hakyimlab/MetaXcan) to be compatible with the MASHR models. We then performed S-MultiXcan analysis of each trait using the MASHR models built off the V8 GTEx release. Associated genes were defined as those found to have a significance of p < 10^-4^ by S-MultiXcan.

#### Global Clustering Coefficient Calculation

The genes identified for an independent GWAS were projected onto the 173 identified clusters. Following this, we generated an unweighted adjacency matrix in which genes in the same cluster were represented by a 1, and genes in different clusters by a 0. A comparison between the same gene was represented by 0. Finally, the global clustering coefficient (GCC) for the genes was calculated. To derive a statistical significance, we randomly sampled the same number of genes as there were significant genes for the phenotype and calculated the GCC one-hundred times. A Z score was calculated from the curve generated by the sampled values.

#### Gene Prediction

We took a simple approach of predicting which clusters were associated with the trait using the S-MultiXcan associations from the smaller GWAS and then checking whether novel gene associations identified by the larger GWAS were in those clusters. A chi-squared enrichment test was used where the minimum expected frequency was greater than 5, and a fisher’s test if not. Several approaches to predict clusters associated with the trait were trialled. The first was to identify any of the 173 clusters with a significant gene in it. The second was to integrate the additional phenotype into the trait-gene association matrix. Next, clusters were identified which had an overall significance signature > 1.5 times the average or were significantly (p < 0.05) higher than the average signature. Different values were tested for these thresholds, with these providing the best performance. The third approach was to predict associated clusters from the previously established 173 clusters using the thresholds taken in approach two. A one-way ANOVA was performed with pairwise comparisons to determine the best approach. Approach three was the most effective, albeit not significantly, while maintaining a low computational burden. In instances where transethnic GWAS were compared, the East-Asian GWAS was used to predict the trait relevant clusters, and the European GWAS was used as the test set.

### Gene Prioritization Analysis

The GWAS with the largest sample size for each of the 13 traits listed in **Table S1** was used to determine the potential of our pipeline for prioritizing genes within a locus. Clumping was performed on each summary statistic using PLINK (https://www.cog-genomics.org/plink/) and 1000 genomes phase 1 genotype data with an LD threshold of 0.5. This was followed by clumping for long distance LD, at the same threshold. Next, we identified individual 500kb regions around the lead SNPs and the genes within that region.

We took a leave one chromosome out (LOCO) approach, where we removed all potential genes on one chromosome. With the remaining genes, we identified which clusters were enriched for genes associated with the trait. To calculate enrichment, we treated all genes associated with one locus as one positive, so that enrichment was for different loci and not genes at the same locus. A fisher’s enrichment test was used to determine significance. Finally, we assessed at what proportion of loci the UnTANGLeD clusters identified a gene when that chromosome was left out of the analysis.

### Normalisation

We trialled relative count, centralised-log ratio and logarithmic normalisation on the chi-squared transformed values of the gene-trait matrix across phenotypes. We evaluated their effects on the following metrics: correlation score, silhouette score, GO and STRING enrichment, global clustering coefficient, prediction of GWAS. A Kruskal Wallis one-way analysis of variance was used to evaluate differences. Relative count was used for the final pipeline.

### Phenotype Filtering Based on Euclidean Distance

A distance matrix between phenotypes using chi-squared transformed, RC-normalised data was generated using the Euclidean distance formula from the package *wordspace*. Phenotypes with a Euclidean distance below a set threshold, which indicated a high degree of relatedness, were removed from the data, leaving the phenotype with the highest number of significant associations. This was performed for thresholds 0 to 62, at which too few phenotypes remained to cluster the genes using the dimensionality reduction methods. GO enrichment was used to evaluate the clustering efficacy at each threshold.

### Phenotype Subsampling and Sensitivity Analyses

Phenotype subsampling was performed on two datasets; MultiXcan results for 1393 phenotypes across 16,849 genes generated in this paper, and another dataset containing MultiXcan results for 4091 phenotypes across 15,734 genes (phenomexcan.org). For the data containing 1393 phenotypes, subsampling was performed randomly without replacement from 50 to 1393 phenotypes in 20 equal increments across 5 replicates for each number of traits. The full UnTANGLeD clustering pipeline was applied to each subsampled matrix. Adjusted rand index (ARI) was calculated for each of the subsampled clustering configurations compared to the full dataset. This analysis was repeated for the data containing 4091 phenotypes; however, subsampling was performed from 50 phenotypes to 4091 phenotypes in 50 equal increments.

### Cluster Conservation

To explore the cause for the marked increase in ARI between 1322 phenotypes and 1393 phenotypes, cluster conservation was calculated between them. For each cluster from 1393 phenotypes, the proportion of genes that remained grouped together in each of the clusters from 1322 phenotypes was calculated. That proportion was used to assign a conservation score to each gene, depending on how large the proportion of cluster the specific gene remained with was. The same approach was applied between 4091 phenotypes and 4009 phenotypes.

## RESULTS

### Unsupervised identification of gene groups with shared complex trait associations

We used MultiXcan results from CTG-VL (10) derived from publicly available GWAS (primarily from UK Biobank, on ∼400,000 individuals) to create a gene-trait association matrix for 16,849 genes and 1,393 traits (Figure 1, Figure S1, Table S2). For each gene trait pair, MultiXcan predicts whether trait-associated variants alter the gene’s expression. The chi-squared transformed significance value for each gene-trait association pair was compiled into the gene-trait association matrix (Figure 2A). These values were normalised using relative count normalisation to account for the difference in power between phenotypes. Performance was not significantly different using other normalisation methods including centralised log ratio or log normalised data (Figure S2). The data was then clustered using Seurat, a dimensionality reduction method commonly used to analyse single cell RNA sequencing data to cluster cells into related groups (11). Here, we use Seurat to test whether the calculated gene-trait associations could be simplified into biologically enriched gene clusters. Clustering was performed across 100 stepwise increases in resolution, a parameter which increases the number of gene clusters. Repeat iterations provided an opportunity to survey both the broad scope of biological processes that could be identified, as well as the specificity that could be achieved with each biological process (Figure 2B).

**Figure 2.**
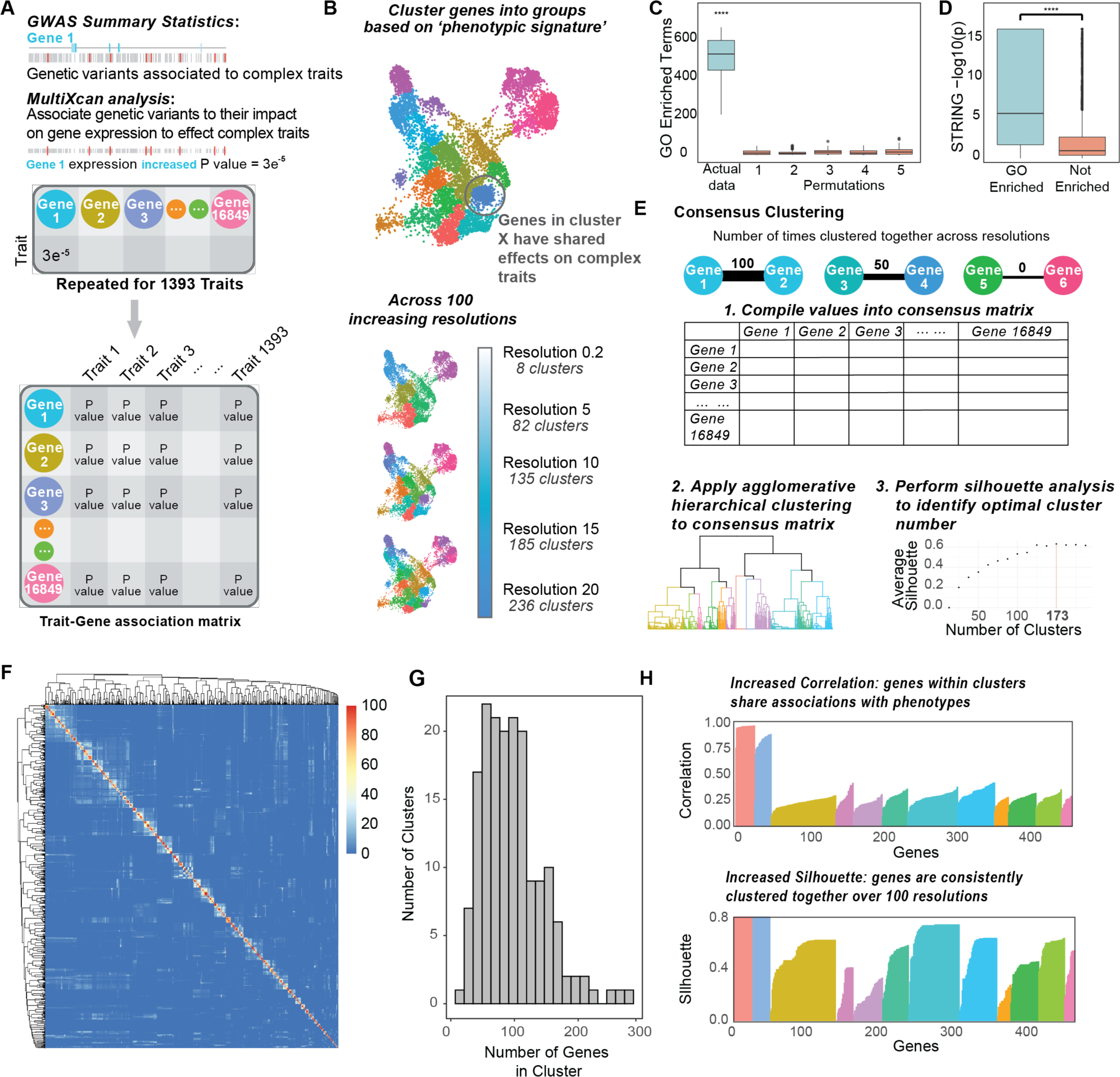
Consensus clustering method identifies biologically enriched gene clusters. **(A)** MultiXcan analysis links genetic variants to genes by predicting changes in gene expression using eQTLs. The chi-squared values of the associations between each of 1393 traits and 25851 genes were compiled into a gene-trait matrix. **(B)** Dimensionality reduction clustering of genes based on their phenotypic associations was performed using *Seurat* across resolutions 0.2 to 20 in 0.2 increments. **(C)** Five permutations of the dataset were compared to the real data by the number of enriched gene ontology terms per resolution. Enrichment was performed using *ClusterProfiler,* FDR corrected p-value < 0.01. Pairwise comparisons between permutations were performed with Wilcoxon signed rank test. **(D)** Validation of gene ontology enriched clusters with STRING protein-protein interaction enrichment. Wilcoxon signed rank test was used to compare STRING enrichment in gene ontology enriched and non-enriched clusters. **(E)** Each gene pair is given a similarity score based on how often they were clustered together across 100 resolutions and these values are compiled into a consensus matrix. Agglomerative hierarchical clustering is applied to the matrix, with the plateau in the average silhouette score defining the optimal number of clusters. **(F)** Heatmap of consensus matrix as clustered using agglomerative hierarchical clustering for 173 clusters. **(G)** Histogram of the number of genes in each of the 173 clusters. **(H)** Silhouette scores and correlation scores calculated for each gene to evaluate the clustering robustness and quality respectively. Data generated for 450 genes selected from 12 random clusters.

To test the biological validity of the derived clusters, we used positive gene sets as defined by gene ontology (GO) (12) and STRING (13) to show that gene clusters have significant enrichment for GO biological processes and STRING protein-protein interactions **(Figure 2C and 2D)**. To demonstrate that the observed enrichment is driven by distinct gene-trait association signatures rather than chance, we performed permutation analyses in which the values in the data matrix were randomly re-ordered. Permutations had significantly fewer enriched clusters, GO terms and a lower strongest significance compared to the real data (p < 4×10^-27^) **(Figure 2C, Figure S3A-B)**. Furthermore, we validated that GO enriched clusters were more likely to also have enrichment for protein-protein interactions, suggesting the enrichment is robust **(Figure 2D)**. This analysis revealed that genes possessing similar associations to complex trait phenotypes cluster meaningfully into biologically enriched groups and the enrichment is not stochastic.

We next developed an ensemble learning method we call “consensus clustering” that incorporates a measure of clustering robustness and quality. Across each of 100 stepwise increases in clustering resolution we evaluated the robustness of clustering by assessing how often every possible gene combination was clustered together ranging from 100 (always) to 0 (never) and compiled these values into a consensus matrix **(Figure 2E)**. Following this, we performed agglomerative hierarchical clustering, evaluating the average silhouette score at each possible number of clusters. The silhouette score quantifies how consistent genes within the same cluster are across *Seurat* resolutions. To derive the optimal number of gene clusters, we calculated the plateau point of the average silhouette score, which informs the number of clusters at which point further splitting no longer improves the stability of clustering assignments **(Figure S3C)**. Applying this methodology to gene-trait associations for 16,849 genes, we identified 173 clusters with an average of 97 genes **(Figure 2F, G)**. Across each cluster, we measured the silhouette score, a metric of cluster robustness and the correlation score, a metric of relation across phenotypes, thereby providing two metrics to quantify the quality of clustering **(Figure 2H)**. Collectively, we call this approach UnTANGLeD: Unsupervised Trait Analysis of Networks from Gene Level Data.

### Consensus clustering identifies robust gene groups enriched for known gene sets

We analysed each cluster by reference to curated annotations of gene programs (GO, disease ontology (14)), signalling pathways (KEGG (15)), protein-protein interactions (STRING), and tissue specificity (16) to evaluate the ability of UnTANGLeD to identify distinct, biologically established gene programs in an unsupervised manner **(Figure 3A, Figure S3D, Tables S3-8)**. This analysis revealed significant enrichment of cell biological pathways and networks across gene clusters, with stronger enrichment among clusters with higher silhouette and correlation scores **(Figure 3A)**. We further performed enrichment analysis of the UnTANGLeD clusters using the EnrichR database (17) **(Figure S4A-C, Table S9)**, finding considerable enrichment for disease-associated genes, gene-expression perturbations associated with disease states or drugs and protein domains and families. We note that although many clusters contain multiple members of a protein family (18) the proportion of any one protein family in the cluster is minor **(Figure S4D, Table S10)**.

**Figure 3.**
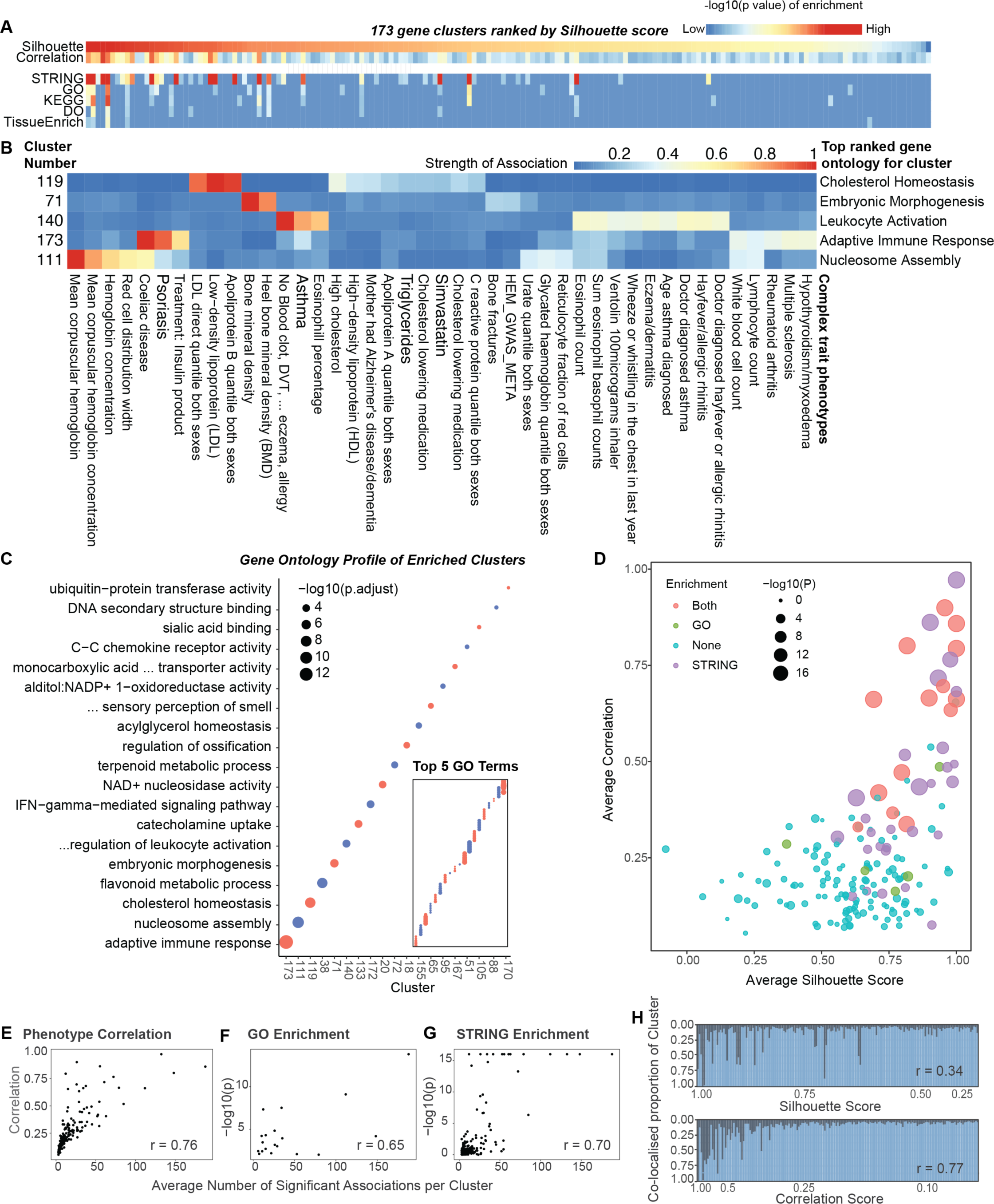
Enrichment of identified clusters for known gene sets is dependent on data quality. **(A)** Broad enrichment profile of 173 clusters stratified by average silhouette and correlation scores. **(B)** Heatmap showing the relationship between the biological profile of five clusters, as proxied by their top gene ontology term, and the unique phenotypic signature. The top 10 phenotypes per cluster were selected. Association strength was calculated using negative log transformed significance values, which were then normalised across phenotypes and then per cluster. **(C)** Top enriched gene ontology biological processes for each cluster have no overlap. Clusters ranked by strength of top enriched term. Specificity of top 5 terms per cluster is also provided. **(D)** 173 clusters stratified by their average correlation and silhouette scores with their gene ontology and STRING enrichment indicated. The P-value represented is specific to the enriched category, and in the case of both represents the more significant of the two. **(E-G)** Correlation between the average number of significant gene-trait associations per cluster and **E)** the average correlation score of each cluster **(F)**, strength of GO enrichment and strength of STRING enrichment per cluster (Pearson’s correlation) **(G)**. **(H)** Presence and degree of colocalization within clusters as stratified by their silhouette and correlation scores (Pearson’s correlation).

We next investigated the relationship between individual gene clusters and the traits most strongly influencing the genes within the clusters, using enriched GO processes as a proxy for the functional profile of a cluster **(Figure 3B)**. Each cluster is defined by a distinct gene-trait association ‘signature’ indicated by the variation and strength of association across 1,393 diverse complex traits. In some instances, the enriched biological processes for certain gene clusters are clearly related to the cluster’s most significantly associated complex trait phenotypes (e.g., cluster 119: GO enrichment: Cholesterol Homeostasis; Dominant complex trait phenotypes: Low-density lipoprotein, Alipoprotein B quantile).

Since UnTANGLeD draws on associations across diverse phenotypes to inform gene-gene relationships, the method can identify gene groups with enriched functions that are apparently biologically independent of the phenotypes most significantly associated with the genes in the cluster. For example, cluster 80, enriched for embryonic morphogenesis (GO:004859), is most significantly associated to the phenotype Bone Mineral Density and cluster 111, enriched for nucleosome organisation (GO:0034728), is most significantly associated to the phenotype Mean Corpuscular Haemoglobin. These results support the central hypothesis that genes with shared effects across diverse phenotypes can be clustered into gene groups controlling shared biological functions and processes in an unsupervised manner **(Figure 3B)**.

Importantly, we show that the GO enriched gene clusters show no overlap in their strongest enriched biological functions, and almost no overlap in their top 5 enriched terms, demonstrating the use of gene-trait association data to parse novel biological gene programs encoded within the genome **(Figure 3C)**.

Stratifying clusters by their silhouette and correlation scores reveals a higher level of GO, STRING, KEGG, DO and tissue specificity enrichment with higher clustering quality, indicating that the metrics provide an accurate representation of cluster quality **(Figure 3A, D)**. Furthermore, both the robustness of clustering and the presence and strength of GO and STRING enrichment are correlated with the number of significant associations to phenotypes per gene (Pearson’s correlation, r > 0.65), as well as the stability of clustering (Pearson’s correlation, r > 0.69) **(Figure 3E-G, Figure S5A-F)**.

Lastly, we note that there is considerable colocalization of genes within clusters, with a stronger relationship between the correlation score and the degree of colocalization for the genes in a cluster (Pearson’s correlation, r = 0.77), than the cluster robustness (Pearson’s correlation, r = 0.34) **(Figure 3H, Table S11)**. STRING enrichment may also be inflated due to the text-mining component, as findings from GWAS may be incorporated into the database, with genes in proximity often being reported together. Indeed, we find that the loss of enrichment due to removal of the text-mining component is correlated with the colocalization of the cluster (r = 0.60) **(Figure S6A-B)**. However, clusters with a high degree of colocalization are not necessarily artefacts of false-positive associations identified by MultiXcan. For example, clusters 173 and 111 are strongly enriched for immune processes and chromatin organisation respectively, despite being highly colocalised **(Figure S6C-D).**

### Subsampling reveals need for more data to improve accuracy of UnTANGLeD

We next sought to determine how the number and diversity of phenotypes influences the accuracy and utility of UnTANGLeD clusters. We show that the number of GO enriched clusters is highly correlated with the number of phenotypes utilised in the analysis (Pearson’s correlation, r = 0.85), even when phenotypic diversity is preserved **(Figure S7A)**. To further test this, we performed phenotype subsampling and evaluated clustering accuracy using an adjusted rand index (ARI) analysis. We found that clustering accuracy compared to the full data improved with the addition of more phenotypes, however a marked increase in ARI between 1322 and the full data set suggests that inaccuracy in clustering that isn’t determined by phenotypic diversity can be attributed to genes which have weak signatures and few significant associations **(Figure S7B)**. We repeated subsampling in a larger dataset containing MultiXcan analysis of 4091 phenotypes retrieved from Pividori *et al*. (2020) which resulted in the same outcome **(Figure S7C)**. Comparison of the two data sets revealed that genes already having many significant associations simply had more associations in the larger dataset with both datasets possessing an equal proportion of genes with few to no significant associations **(Figure S7D)**. Further, that genes with higher numbers of significant associations have higher degrees of conservation **(Figure S7E-F)**. It’s likely that the effective number of traits is similar between the two datasets, as both mostly draw on the UK Biobank and have many highly correlated phenotypes

Cumulatively, these findings indicate that the quality of gene clustering is dependent on the scale and quality of data needed to derive high silhouette and correlation scores as a basis for efficient enrichment of functional gene clusters. Accordingly, as more data becomes available, the quality and accuracy of UnTANGLeD will improve. However, simply increasing the number of phenotypes leads to an increase in redundant associations, and therefore strategies to increase the number of significant gene-trait associations across the genome should be employed, such as diversifying phenotypes and increasing cohort size.

### UnTANGLeD clusters are conserved across traits and can predict novel trait associated genes

GWAS require collections of large cohorts comprising thousands of individual-level genotype data to characterise the genetic architecture of a trait. Furthermore, collecting enough samples can prove challenging for many diseases, and as such they are often underrepresented in biobanks.

We hypothesised that UnTANGLeD gene clusters would be conserved across complex traits. To test this, we investigated an independent GWAS of ulcerative colitis (UC) (19) **(Figure 4A)**. We show that the 278 genes associated with UC (p < 10^-4^) **(Figure 4B)** were significantly more clustered within the UnTANGLeD clusters than expected by chance (p = 2×10^-9^) **(Figure 4C)**. The result shows that despite not being used to construct the clusters, UC associated genes nevertheless group within the UnTANGLeD clusters, demonstrating that the defined gene programs are conserved. We replicate our findings in 6 additional independent GWAS phenotypes, highlighting that the UnTANGLeD clusters are conserved across a broad phenotypic space (3, 20–28) **(Figure 4G)**.

**Figure 4.**
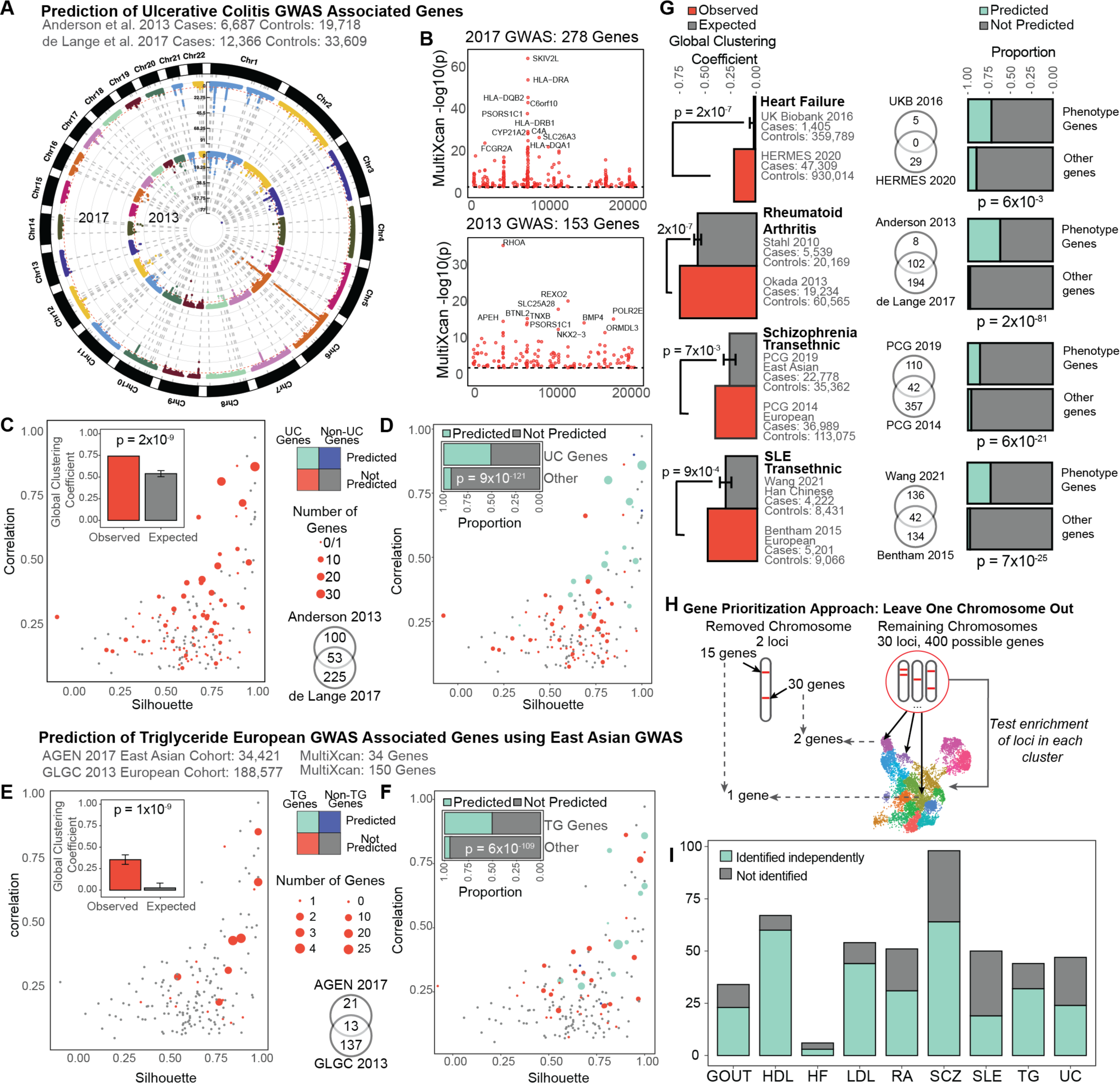
Identified clusters are conserved across all phenotypes and can be used for prediction of genes involved in complex trait biology and prioritization of GWAS genes at implicated loci. **(A)** Manhattan plot of loci identified by a 2017 and 2013 GWAS of ulcerative colitis (UC). **(B)** Manhattan plot of S-MultiXcan genes for the two GWAS respectively, genes are ordered according to their genomic positions. **(C)** Distribution of 278 significant genes from the 2017 UC GWAS across 173 clusters. Global clustering coefficient was calculated for the 278 genes. Significance was calculated using 100 bootstrap replicates to establish a distribution from which a Z score was calculated. **(D)** Prediction of 2017 UC GWAS genes using 2013 UC GWAS. Chi-squared enrichment test was used to determine enrichment for prediction of novel genes compared to non-trait associated genes. **(E)** Distribution and global clustering coefficient of 34 significant genes from East Asian GWAS of Triglyceride levels. Significance was calculated using bootstrapping. **(F)** Prediction of 137 novel genes from 2013 European GWAS of triglycerides using 2017 East Asian GWAS. Enrichment was calculated using chi-squared test. **(G)** Increase in observed global clustering coefficient compared to expected and prediction enrichment across four additional traits. **(H)** Schematic of gene prioritization strategy using the leave one chromosome out approach. Potential genes at a significant locus were refined using clusters enriched for the trait. **(I)** The proportion of loci at which a gene was successfully identified independently of all genes on the same chromosome.

We next tested whether the gene clusters can be used to predict novel genes and cellular processes underpinning independent complex trait data. To test this hypothesis, we examined two GWAS for UC. The first was performed in 2013 with 6,687 cases and 19,718 controls (29), and the latter in 2017 with 12,366 cases and 33,609 controls (19) **(Figure 4A)**. MultiXcan analysis of the summary statistics identified 153 and 278 genes respectively, with an overlap of 53 genes **(Figure 4B)**. We projected the MultiXcan associations for the 2013 GWAS onto the 173 clusters, identifying clusters were statistically associated with UC **(Figure S8)**. Finally, we tested whether the clusters predicted from the 2013 GWAS contained novel genes identified by the 2017 GWAS. Of the 225 novel genes identified by the 2017 GWAS, our approach was able to use the 2013 GWAS to predict 120 with a significant enrichment for predicting UC associated genes compared to other genes (p < 3×10^-121^, chi-squared test) **(Figure 4D)**.

GWAS of the same complex trait conducted in populations of differing ancestries may implicate both shared and distinct loci. We tested whether UnTANGLeD clusters are conserved for genes specific to non-European ancestries, given that the UnTANGLeD gene clusters are built from a European cohort. To test this, we examined a GWAS for triglyceride levels in an East Asian population, which identified 34 genes (30) **(Figure S9A-B)**. Mirroring our findings in a GWAS conducted on a European population, we found that the genes associated with triglyceride levels in an East Asian population are significantly more clustered than expected (p = 1×10^-9^) and replicate this finding in 4 other GWAS conducted in populations of non-European ancestry (30–32). We further tested whether the GWAS conducted in the East Asian cohort could be used to predict novel genes identified in a European cohort. We found that clusters implicated in triglyceride levels using the East Asian GWAS were highly enriched for genes identified by the European GWAS (p = 6×10^-109^) **(Figure 4F)**.

All together, we show significant enrichment for prediction of novel genes across GWAS performed for 7 traits in differing cohort sizes in a European population, and 4 traits for which GWAS were performed in different ancestries (3, 20-28, 30–33) **(Figure 4G, Figure S9C)**.

We further tested whether the UnTANGLeD clusters could be used to prioritize causal genes at any given locus. It is difficult to accurately identify the causal genes from GWAS identified variants due to linkage disequilibrium and complex regulatory effects of intergenic variants. For each independent trait, we identified potential gene candidates within 500kb of each independent significant SNP then took a leave one chromosome out approach (LOCO) to investigate whether genes on the removed chromosome would be implicated in the clusters associated with the remaining genes. **(Figure 4H)**. We are able to identify a major proportion of loci independently across all traits and reduce the potential candidates at each locus considerably, further highlighting the utility of UnTANGLeD **(Figure 4I)**.

## DISCUSSION

This study demonstrates that gene programs governing biological processes can be identified without reliance on prior knowledge, by analysing the association between genetic variation and a large range of diverse complex traits. Several prior studies have constructed small gene networks using a limited number of disease phenotypes and their associated genes from curated GWAS databases and restricted sources of rare genetic variants. Other studies, like PheWAS (34, 35) and PhenomeXcan (36) have collated genomic associations across numerous phenotypes to create resources of variant-trait and gene-trait associations.

Here, we construct a gene-trait association matrix for 16,849 genes across 1,393 complex traits similarly to PhenomeXcan, and further the concept by using UnTANGLeD to identify gene programs. We apply dimensionality reduction methods, which can harness the high dimensional, complex gene-trait association data, allowing us to greatly expand on the scale of studies previously attempting to build gene networks. By increasing the scale of data, we not only identify gene programs enriched for biological processes specific to associated phenotypes but also reveal gene programs enriched for central processes governing diverse mechanisms of cellular development and homeostasis.

The UnTANGLeD framework is a powerful approach to identify gene programs orchestrating key biological processes. We implicate novel genes in clusters enriched for known processes and identify numerous novel gene programs with enrichment for protein-protein interactions and no known function. We further highlight the utility of UnTANGLeD for hypothesis generation and functional annotation of genes, which may be particularly valuable for non-coding genes, as they are notoriously difficult to annotate *in silico* (37). Finally, the UnTANGLeD framework reveals relationships between complex traits, linking phenotypes by the gene programs that underpin them.

We demonstrate the utility of UnTANGLeD for predicting genes associated with complex traits and diseases using a low-powered GWAS of the same trait. Currently, standard methods use gene-set analysis to improve power to identify genes and pathways involved in a phenotype, such as MAGMA, or GIGSEA (38–41). Our method eliminates the need to define gene-sets and instead uses gene-trait association data to learn gene sets governing complex traits (39), enabling us to implicate novel trait associated genes and loci from a much smaller cohort size.

We further highlight the use of the UnTANGLeD clusters for gene prioritization, showing that they effectively select gene candidates at different loci related to the same phenotype. Current gene prioritisation approaches use either distance-based metrics or mapping to eQTLs to predict changes in gene expression (42). However, these also suffer from a considerable false positive rate and may not always distinguish between two genes in proximity, as noted in our data (42). Some recent methods have integrated biological data, such as gene sets, RNA sequencing and protein-protein interaction databases to further prioritise genes at a locus (43). Our framework can be used independently or integrated with any of these approaches to advance understanding of complex trait biology.

Outside of its utility in GWAS analyses, UnTANGLeD may provide key mechanisms for data analysis in medical and industry pipelines including genetic testing and drug discovery. For example, polygenic risk scores (PRS) are an emerging method that evaluate an individual’s disease risk from genetic variants (44). Methods such as UnTANGLeD may help reveal genes and hence genetic variants governing cell programs underlying disease risk and hence improve prediction accuracy. In the context of pharmacogenomics, studies have shown that drug targets with genetic support from either rare or common diseases are more than twice as likely to pass through clinical trials (45, 46). Since UnTANGLeD captures gene programs associated with all complex traits and diseases, its predictive power may help de-risk candidates and thereby decrease cost associated with the drug discovery pipeline. Overall, UnTANGLeD represents a powerful and versatile framework for studying cellular gene programs to interpret diverse sources of orthogonal genetic data.

We note several limitations in our method. Primarily, that the current GWAS data does not represent the whole phenome. Furthermore, many traits are highly correlated, and disease traits are underrepresented in the UK Biobank, the main source of data in this study. Secondly, UnTANGLeD relies on S-MultiXcan to construct the gene-trait association matrix. While S-MultiXcan is powered to detect associations across all tissues, it suffers from a high false discovery rate and may perform poorly in tissues with small sample sizes. Moreover, S-MultiXcan can identify genes colocalised with a causal gene as significant, which can obscure biological signatures. Other approaches such as SMR MR-JTI may remedy this issue (47). Additionally, UnTANGLeD does not account for the predicted directionality of effect or tissue-specific effects, which may help to further increase the quality and biological specificity of the clusters. Biological validation of the method using established gene sets may be inflated due to

GWAS data being included in the definition of the gene sets. Finally, we note that although UnTANGLeD is a powerful tool for identifying clusters in an unsupervised manner, the overall function of the cluster may be difficult to determine. The development of improved gene-based tests and emergence of larger GWAS data spanning the whole phenome will improve the accuracy and utility of UnTANGLeD.

This study provides a powerful framework for the identification of gene programs governing biological processes conserved across all complex traits and diseases, with important applications for functional annotation, hypothesis generation, machine learning and prediction algorithms and interpretation of GWAS and diverse other genomic data types. Our approach can be applied to any collection of gene-trait information, harnessing the power of biological patterns in a diverse landscape of phenotypic variation.

## DATA AND MATERIALS AVAILABILITY

All source code available on GitHub (https://github.com/palpant-comp/UnTANGLeD) and all data available on Zenodo (https://doi.org/10.5281/zenodo.6572617).

## SUPPLEMENTARY MATERIALS

Supplementary Data are available at NAR online.

## FUNDING

We thank support from the National Health and Medical Research Council of Australia (Grant 1143163, N.P.) and the Australian Research Council (Grant SR1101002, N.P.) and the National Heart Foundation of Australia (Grant 101889, N.P.).

## ACKNOWLEDGEMENTS

We thank Naomi Wray and Jian Yang for assistance in conceptual development of the study.

## AUTHOR CONTRIBUTIONS

**DM:** Developed the study, performed all analyses, and wrote the manuscript

**MS** and **CN:** Helped supervise bioinformatics analysis

**GCP:** Helped supervise and design GWAS data selection and analysis, interpreted data, and wrote the manuscript

**NP:** Conceived and supervised the project, raised funding, and wrote the manuscript

## CONFLICT OF INTEREST STATEMENT

GCP is currently an employee of 23andMe Inc. and holds stock options for the company.

## Notes

### Competing Interest Statement

Gabriel Cuellar Partida is currently an employee of 23andMe Inc. and holds stock options for the company.

## REFERENCE

1. ENCODE Project Consortium (2012) An integrated encyclopedia of DNA elements in the human genome. Nature, 489, 57–74.

2. Rozenblatt-Rosen,O., Stubbington,M.J.T., Regev,A. and Teichmann,S.A. (2017) The Human Cell Atlas: from vision to reality. Nat. 2017 5507677, 550, 451–453.

3. Sudlow,C., Gallacher,J., Allen,N., Beral,V., Burton,P., Danesh,J., Downey,P., Elliott,P., Green,J., Landray,M., et al. (2015) UK Biobank: An Open Access Resource for Identifying the Causes of a Wide Range of Complex Diseases of Middle and Old Age. PLoS Med., 12, 1–10.

4. Jumper,J., Evans,R., Pritzel,A., Green,T., Figurnov,M., Ronneberger,O., Tunyasuvunakool,K., Bates,R., Žídek,A., Potapenko,A., et al. (2021) Highly accurate protein structure prediction with AlphaFold. Nature, 596, 583–589.

5. Shim,W.J., Sinniah,E., Xu,J., Vitrinel,B., Alexanian,M., Andreoletti,G., Shen,S., Sun,Y., Balderson,B., Boix,C., et al. (2020) Conserved Epigenetic Regulatory Logic Infers Genes Governing Cell Identity. Cell Syst, 11, 625–639.e13.

6. Visscher,P.M., Wray,N.R., Zhang,Q., Sklar,P., McCarthy,M.I., Brown,M.A. and Yang,J. (2017) 10 Years of GWAS Discovery: Biology, Function, and Translation. Am. J. Hum. Genet., 101, 5–22.

7. Bellomo,T.R., Bone,W.P., Chen,B.Y., Gawronski,K.A.B., Zhang,D., Park,J., Levin,M., Tsao,N., Klarin,D., Lynch,J., et al. (2021) Multi-trait GWAS of atherosclerosis detects novel pleiotropic loci. medRxiv, 10.1101/2021.05.21.21257493.

8. Yu,G., Wang,L.-G., Han,Y., He,Q.-Y. (2012) clusterProfiler: an R Package for Comparing Biological Themes Among Gene Clusters. Omi. A J. Integr. Biol., 16, 284–287.

9. Liberzon,A., Subramanian,A., Pinchback,R., Thorvaldsdóttir,H., Tamayo,P. and Mesirov,J.P. (2011) Molecular signatures database (MSigDB) 3.0. Bioinformatics, 27, 1739.

10. Cuellar-Partida,G., Lundberg,M., Kho,P.F., D’Urso,S., Gutierrez-Mondragon,L.F. and Hwang,L.-D. (2019) Complex-Traits Genetics Virtual Lab: A community-driven web platform for post-GWAS analyses. bioRxiv, 10.1101/518027.

11. Butler,A., Hoffman,P., Smibert,P., Papalexi,E. and Satija,R. (2018) Integrating single-cell transcriptomic data across different conditions, technologies, and species. Nat. Biotechnol., 36, 411–420.

12. Ashburner,M., Ball,C.A., Blake,J.A., Botstein,D., Butler,H., Cherry,J.M., Davis,A.P., Dolinski,K., Dwight,S.S., Eppig,J.T., et al. (2000) Gene ontology: tool for the unification of biology. The Gene Ontology Consortium. Nat. Genet., 25, 25–29.

13. Szklarczyk,D., Gable,A.L., Lyon,D., Junge,A., Wyder,S., Huerta-Cepas,J., Simonovic,M., Doncheva,N.T., Morris,J.H., Bork,P., et al. (2019) STRING v11: protein–protein association networks with increased coverage, supporting functional discovery in genome-wide experimental datasets. Nucleic Acids Res., 47, D607–D613.

14. Schrimi,L.M., Arze,C., Nadendla,S., Wayne Chang,Y.-W., Mazaitis,M., Felix,V., Feng,G. and Kibbe,W.A. (2012) Disease Ontology: a backbone for disease semantic integration. Nucleic Acids Res., 40.

15. Kanehisa,M. (2000) KEGG: Kyoto Encyclopedia of Genes and Genomes. Nucleic Acids Res., 28, 27–30.

16. Jain,A. and Tuteja,G. (2019) TissueEnrich: Tissue-specific gene enrichment analysis. Bioinformatics, 35, 1966–1967.

17. Kuleshov,M. V, Jones,M.R., Rouillard,A.D., Fernandez,N.F., Duan,Q., Wang,Z., Koplev,S., Jenkins,S.L., Jagodnik,K.M., Lachmann,A., et al. (2016) Enrichr: a comprehensive gene set enrichment analysis web server 2016 update. Nucleic Acids Res., 44, W90–W97.

18. El-Gebali,S., Mistry,J., Bateman,A., Eddy,S.R., Luciani,A., Potter,S.C., Qureshi,M., Richardson,L.J., Salazar,G.A., Smart,A., et al. (2019) The Pfam protein families database in 2019. Nucleic Acids Res., 47, D427–D432.

19. de Lange,K.M., Moutsianas,L., Lee,J.C., Lamb,C.A., Luo,Y., Kennedy,N.A., Jostins,L., Rice,D.L., Gutierrez-Achury,J., Ji,S.G., et al. (2017) Genome-wide association study implicates immune activation of multiple integrin genes in inflammatory bowel disease. Nat Genet, 49, 256–261.

20. Köttgen,A., Albrecht,E., Teumer,A., Vitart,V., Krumsiek,J., Hundertmark,C., Pistis,G., Ruggiero,D., O’Seaghdha,C.M., Haller,T., et al. (2013) Genome-wide association analyses identify 18 new loci associated with serum urate concentrations. Nat Genet, 45, 145–154.

21. Tin,A., Marten,J., Halperin Kuhns,V.L., Li,Y., Wuttke,M., Kirsten,H., Sieber,K.B., Qiu,C., Gorski,M., Yu,Z., et al. (2019) Target genes, variants, tissues and transcriptional pathways influencing human serum urate levels. Nat. Genet., 51, 1459–1474.

22. Shah,S., Henry,A., Roselli,C., Lin,H., Sveinbjörnsson,G., Fatemifar,G., Hedman,Å.K., Wilk,J.B., Morley,M.P., Chaffin,M.D., et al. (2020) Genome-wide association and Mendelian randomisation analysis provide insights into the pathogenesis of heart failure. Nat. Commun., 11, 163.

23. Kathiresan,S., Melander,O., Guiducci,C., Surti,A., Burtt,N.P., Rieder,M.J., Cooper,G.M., Roos,C., Voight,B.F., Havulinna,A.S., et al. (2008) Six new loci associated with blood low- density lipoprotein cholesterol, high-density lipoprotein cholesterol or triglycerides in humans. Nat Genet, 40, 189–197.

24. Willer,C.J., Schmidt,E.M., Sengupta,S., Peloso,G.M., Gustafsson,S., Kanoni,S., Ganna,A., Chen,J., Buchkovich,M.L., Mora,S., et al. (2013) Discovery and refinement of loci associated with lipid levels. Nat. Genet., 45, 1274–1283.

25. Stahl,E.A., Raychaudhuri,S., Remmers,E.F., Xie,G., Eyre,S., Thomson,B.P., Li,Y., Kurreeman,F.A., Zhernakova,A., Hinks,A., et al. (2010) Genome-wide association study meta-analysis identifies seven new rheumatoid arthritis risk loci. Nat Genet, 42, 508– 514.

26. Okada,Y., Wu,D., Trynka,G., Raj,T., Terao,C., Ikari,K., Kochi,Y., Ohmura,K., Suzuki,A., Yoshida,S., et al. (2014) Genetics of rheumatoid arthritis contributes to biology and drug discovery. Nature, 506, 376–381.

27. Schizophrenia Psychiatric Genome-Wide Association Study (GWAS)Consortium (2011) Genome-wide association study identifies five new schizophrenia loci. Nat Genet, 43, 969–976.

28. Schizophrenia Working Group of the Psychiatric Genomics Consortium (2014) Biological insights from 108 schizophrenia-associated genetic loci. Nature, 511, 421–427.

29. Anderson,C.A., Boucher,G., Lees,C.W., Franke,A., D’Amato,M., Taylor,K.D., Lee,J.C., Goyette,P., Imielinski,M., Latiano,A., et al. (2011) Meta-analysis identifies 29 additional ulcerative colitis risk loci, increasing the number of confirmed associations to 47. Nat Genet, 43, 246–252.

30. Spracklen,C.N., Chen,P., Kim,Y.J., Wang,X., Cai,H., Li,S., Long,J., Wu,Y., Wang,Y.X., Takeuchi,F., et al. (2017) Association analyses of East Asian individuals and transancestry analyses with European individuals reveal new loci associated with cholesterol and triglyceride levels. Hum Mol Genet, 26, 1770–1784.

31. Lam,M., Chen,C.-Y., Li,Z., Martin,A.R., Bryois,J., Ma,X., Gaspar,H., Ikeda,M., Benyamin,B., Brown,B.C., et al. (2019) Comparative genetic architectures of schizophrenia in East Asian and European populations. Nat. Genet., 51, 1670–1678.

32. Wang,Y.-F., Zhang,Y., Lin,Z., Zhang,H., Wang,T.-Y., Cao,Y., Morris,D.L., Sheng,Y., Yin,X., Zhong,S.-L., et al. (2021) Identification of 38 novel loci for systemic lupus erythematosus and genetic heterogeneity between ancestral groups. Nat. Commun., 12, 772.

33. Bentham,J., Morris,D.L., Graham,D.S.C., Pinder,C.L., Tombleson,P., Behrens,T.W., Martín,J., Fairfax,B.P., Knight,J.C., Chen,L., et al. (2015) Genetic association analyses implicate aberrant regulation of innate and adaptive immunity genes in the pathogenesis of systemic lupus erythematosus. Nat Genet, 47, 1457–1464.

34. Diogo,D., Tian,C., Franklin,C.S., Alanne-Kinnunen,M., March,M., Spencer,C.C.A., Vangjeli,C., Weale,M.E., Mattsson,H., Kilpeläinen,E., et al. (2018) Phenome-wide association studies across large population cohorts support drug target validation. Nat. Commun. 2018 91, 9, 1–13.

35. Pendergrass,S.A., Buyske,S., Jeff,J.M., Frase,A., Dudek,S., Bradford,Y., Ambite,J.-L., Avery,C.L., Buzkova,P., Deelman,E., et al. (2019) A phenome-wide association study (PheWAS) in the Population Architecture using Genomics and Epidemiology (PAGE) study reveals potential pleiotropy in African Americans. PLoS One, 14, e0226771.

36. Pividori,M., Rajagopal,P.S., Barbeira,A., Liang,Y., Melia,O., Bastarache,L., Park,Y., Consortium,G., Wen,X. and Im,H.K. (2020) PhenomeXcan: Mapping the genome to the phenome through the transcriptome. Sci. Adv., 6, eaba2083.

37. Perron,U., Provero,P. and Molineris,I. (2017) In silico prediction of lncRNA function using tissue specific and evolutionary conserved expression. BMC Bioinformatics, 18, 29–39.

38. Zhu,S., Qian,T., Hoshida,Y., Shen,Y., Yu,J. and Hao,K. (2019) GIGSEA: genotype imputed gene set enrichment analysis using GWAS summary level data. Bioinformatics, 35, 160–163.

39. Zhu,X. and Stephens,M. (2018) Large-scale genome-wide enrichment analyses identify new trait-associated genes and pathways across 31 human phenotypes. Nat. Commun. (2018) 91, 9, 1–14.

40. de Leeuw,C.A., Mooij,J.M., Heskes,T. and Posthuma,D. (2015) MAGMA: Generalized Gene-Set Analysis of GWAS Data. PLOS Comput. Biol., 11, e1004219.

41. Sun,R., Hui,S., Bader,G.D., Lin,X. and Kraft,P. (2019) Powerful gene set analysis in GWAS with the Generalized Berk-Jones statistic. PLoS Genet., 15.

42. Broekema,R. V., Bakker,O.B. and Jonkers,I.H. (2020) A practical view of fine-mapping and gene prioritization in the post-genome-wide association era. Open Biol., 10.

43. Schaid,D.J., Chen,W. and Larson,N.B. (2018) From genome-wide associations to candidate causal variants by statistical fine-mapping. Nat. Rev. Genet., 19, 491.

44. Lewis,C.M. and Vassos,E. (2020) Polygenic risk scores: From research tools to clinical instruments. Genome Med., 12, 1–11.

45. Nelson,M.R., Tipney,H., Painter,J.L., Shen,J., Nicoletti,P., Shen,Y., Floratos,A., Sham,P.C., Li,M.J., Wang,J., et al. (2015) The support of human genetic evidence for approved drug indications. Nat. Genet. 2015 478, 47, 856–860.

46. King,E.A., Wade Davis,J. and Degner,J.F. (2019) Are drug targets with genetic support twice as likely to be approved? Revised estimates of the impact of genetic support for drug mechanisms on the probability of drug approval. PLOS Genet., 15, e1008489.

47. Zhou,D., Jiang,Y., Zhong,X., Cox,N.J., Liu,C. and Gamazon,E.R. (2020) A unified framework for joint-tissue transcriptome-wide association and Mendelian randomization analysis. Nat. Genet., 52, 1239–1246.

