## Supplementary Materials for "Organisation of gene programs revealed by unsupervised analysis of diverse gene-trait associations"

[#] Co-corresponding authors

*Current address: 23andMe Inc.

**Corresponding authors**

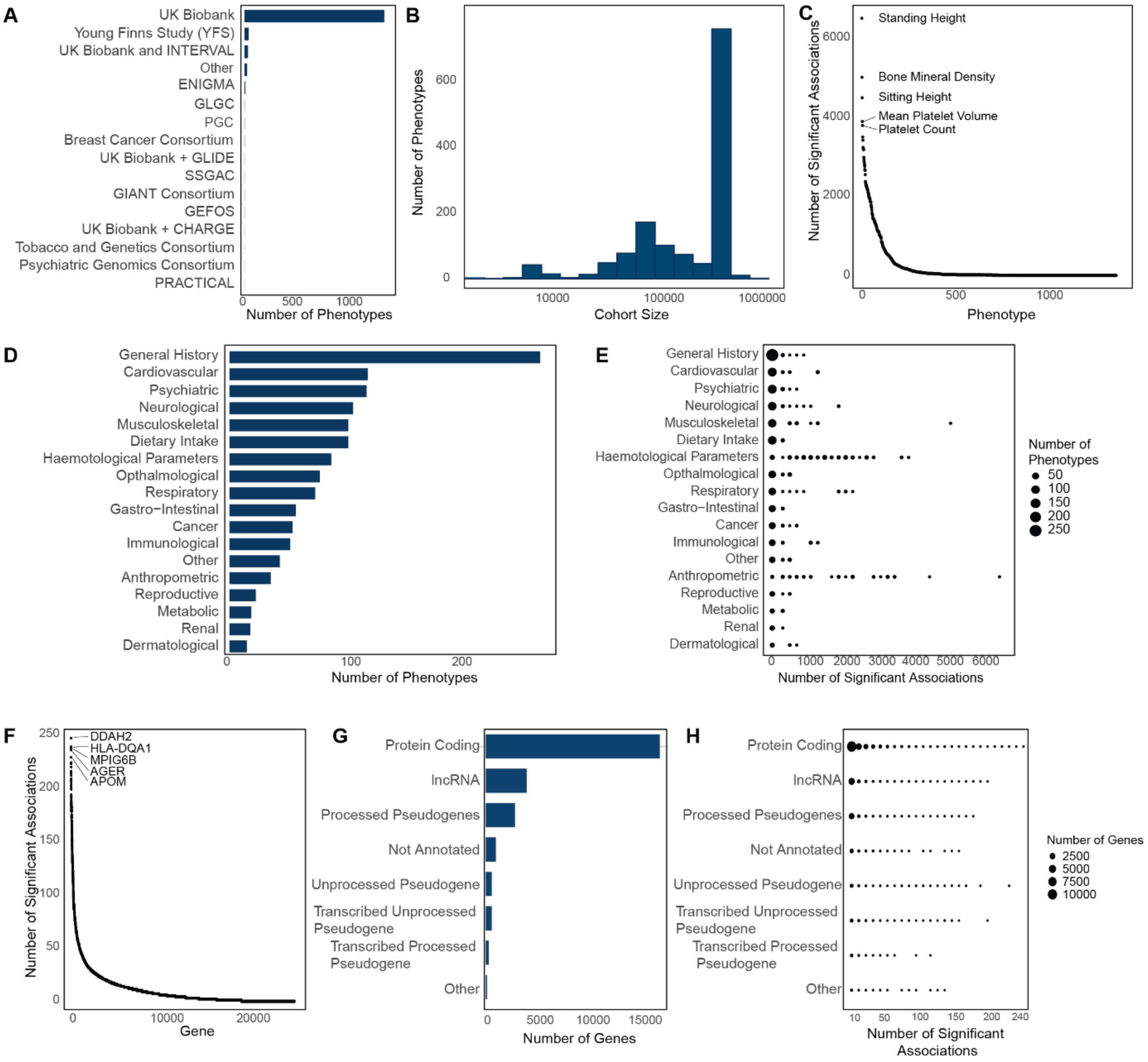


**Supplementary Figure 1. Breakdown of GWAS cohorts, cohort sizes and gene-phenotype associations.**

**(A)** Source consortium for GWAS summary statistics with their respective contributions.

**(B)** Total cohort size for each phenotype, shown on a logarithmic scale.

**(C)** The number of significant associations ($p < 10^{-4}$) identified by MultiXcan per phenotype. The top 5 phenotypes have been annotated.

**(D)** Categorization of phenotypes within the data by the physiological system affected.

**(E)** The number of significant associations per phenotype, using the same categorization as in D.

**(F)** The number of significant associations identified by MultiXcan per gene.

**(G)** Categorization of genes by *biomaRt* gene biotype. 1000 genes were not recognised.

**(H)** The number of significant associations per gene categorized by gene biotype.

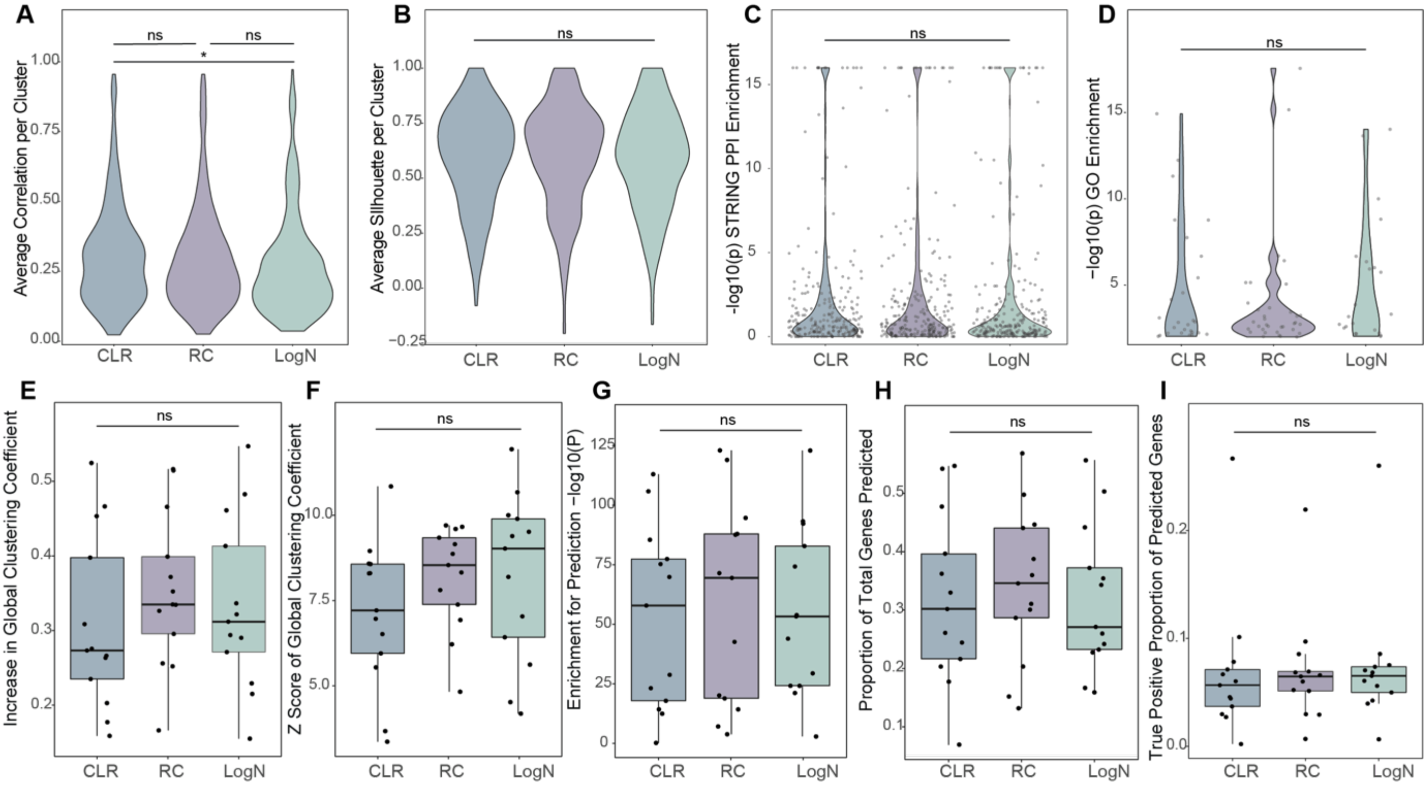


**Supplementary Figure 2. Comparison of normalisation methods on performance.**

(* = P < 0.05)

**(A-D)** Distribution of Correlation scores **(A)**, Silhouette scores **(B)**, STRING enrichment values **(C)** and GO enrichment values **(D)** when significance values were normalised using centralised log ratio (CLR), relative count (RC) or log normalisation. A Kruskal Wallis one way analysis of variance test was used, followed by the Dunn test for pairwise comparisons.

**(E-F)** Effect of normalisation method on observed increase in global clustering coefficient **(E)** and Z score of increase of genes associated with 13 independent phenotypes **(F)**.

**(G-I)** Effect of normalisation method on enrichment for prediction of genes **(G)**, proportion of genes predicted **(H)**, and proportion of predictions that are correct **(I)**.

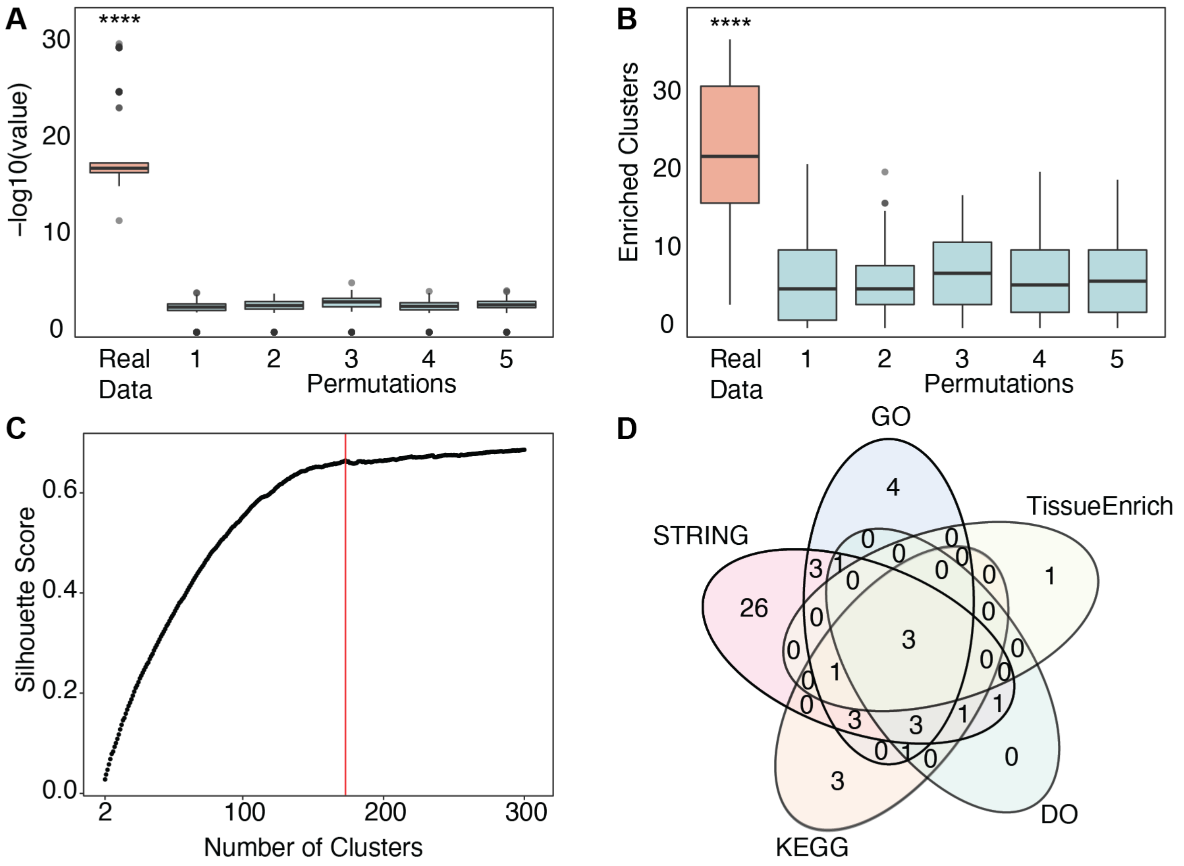


**Supplementary Figure 3. Exploratory analysis of clustering genes using their phenotypic signature.**

**(A-B)** Five permutations of the dataset were compared to the real data based on the strongest GO enrichment **(A)** and the number of enriched clusters per resolution **(B)**. The Wilcoxon test was performed and corrected for multiple testing. (**** P < 0.0001)

**(C)** Average silhouette score calculated from 2 to 300 clusters for hierarchical clustering performed on the consensus matrix. The optimal number of clusters was determined at the inflection point of 173 clusters.

**(D)** Venn diagram of broad enrichment analysis across 173 clusters as performed using *ClusterProfiler* for disease ontology (DO), gene ontology (GO), KEGG pathways, STRING protein-protein interactions and tissue specificity (TissueEnrich).

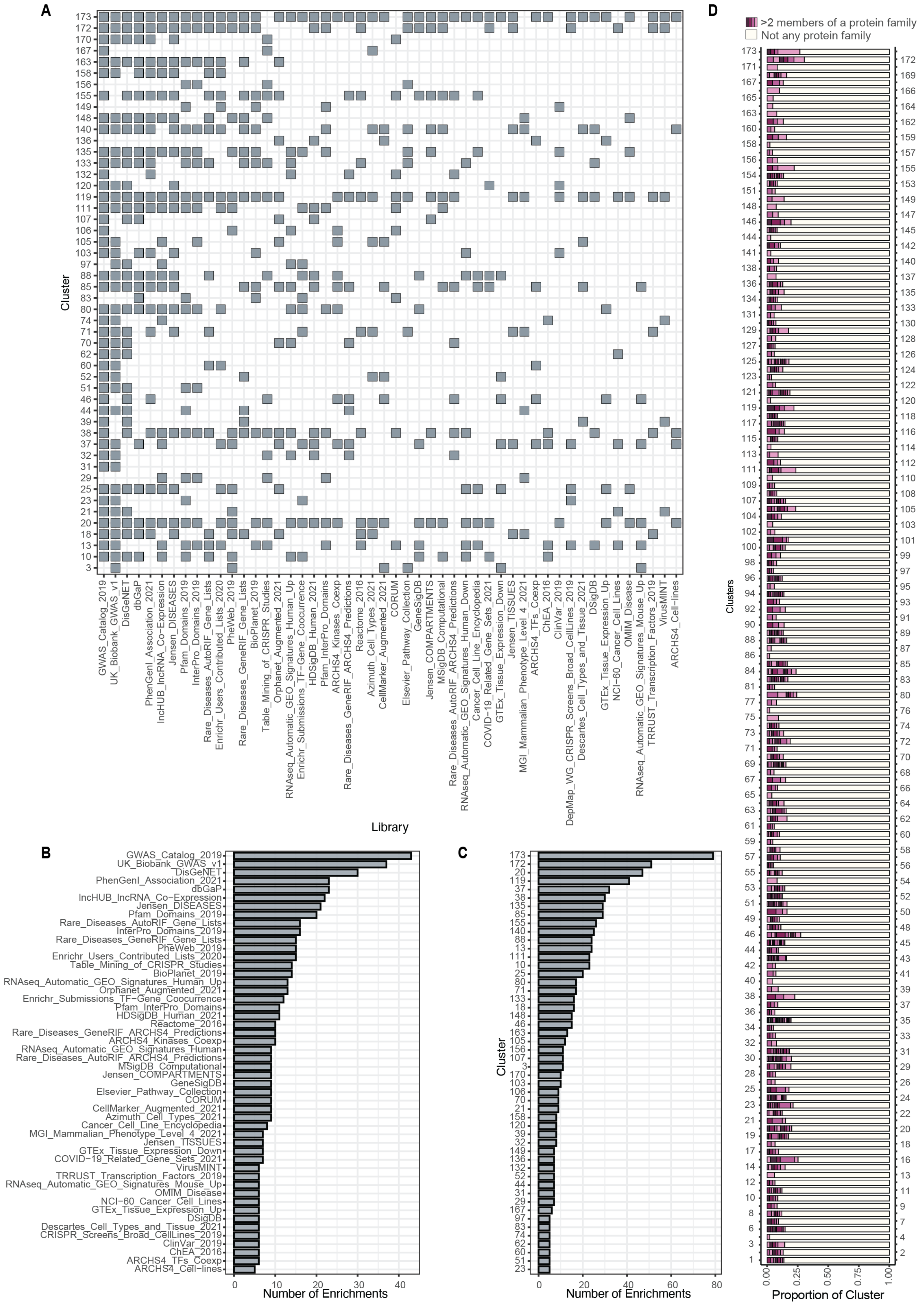
A
Cluster
Library
B
Number of Enrichments
GWAS_Catalog_2019
UK_Biobank_GWAS_v1
DisGeNET
PhenGenI_Association_2021
dbGaP
lncHUB_lncRNA_Co−Expression
Jensen_DISEASES
Pfam_Domains_2019
Rare_Diseases_AutoRIF_Gene_Lists
InterPro_Domains_2019
Rare_Diseases_GeneRIF_Gene_Lists
PheWeb_2019
Enrichr_Users_Contributed_Lists_2020
Table_Mining_of_CRISPR_Studies
BioPlanet_2019
RNAseq_Automatic_GEO_Signatures_Human_Up
Orphanet_Augmented_2021
Enrichr_Submissions_TF−Gene_Coocurrence
Pfam_InterPro_Domains
HDSigDB_Human_2021
Reactome_2016
Rare_Diseases_GeneRIF_ARCHS4_Predictions
ARCHS4_Kinases_Coexp
RNAseq_Automatic_GEO_Signatures_Human
Rare_Diseases_AutoRIF_ARCHS4_Predictions
MSigDB_Computational
Jensen_COMPARTMENTS
GeneSigDB
Elsevier_Pathway_Collection
CORUM
CellMarker_Augmented_2021
Azimuth_Cell_Types_2021
Cancer_Cell_Line_Encyclopedia
MGI_Mammalian_Phenotype_Level_4_2021
Jensen_TISSUES
GTEx_Tissue_Expression_Down
COVID−19_Related_Gene_Sets_2021
VirusMINT
TRRUST_Transcription_Factors_2019
RNAseq_Automatic_GEO_Signatures_Mouse_Up
OMIM_Disease
NCI−60_Cancer_Cell_Lines
GTEx_Tissue_Expression_Up
DSigDB
Descartes_Cell_Types_and_Tissue_2021
CRISPR_Screens_Broad_CellLines_2019
ClinVar_2019
ChEA_2016
ARCHS4_TFs_Coexp
ARCHS4_Cell−lines
C
Cluster
Number of Enrichments
D
>2 members of a protein family
Not any protein family
Clusters
Proportion of Cluster

**Supplementary Figure 4. Broad Enrichment performed using EnrichR.**

**(A)** Significant enrichment across the top 50 clusters and top 50 EnrichR libraries as ranked by the number of significant enrichments. Gene ontology, KEGG and chromosomal location libraries were excluded due to redundancy with analyses already performed.

(**B, C)** Number of significantly enriched (**B)** Clusters per EnrichR library and (**C)** EnrichR libraries per Cluster.

(**D)** Representation of shared protein domains within clusters as calculated by EnrichR for the Pfam_Domains_2019 library. Differently coloured segments represent proteins belonging to distinct families.

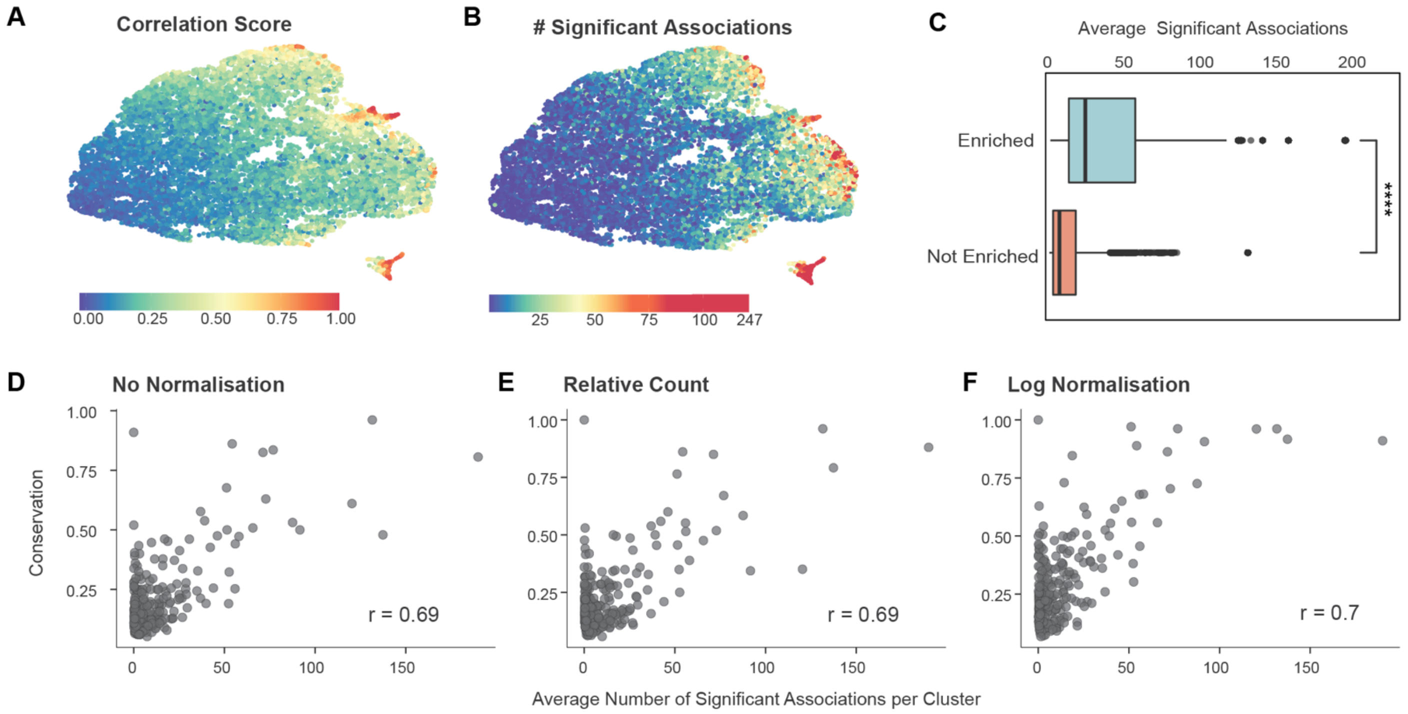


**Supplementary Figure 5. Influence of data scale on quality of clustering.**

**(A-B)** UMAP of 16,849 genes generated by *Seurat* coloured by correlation with neighbouring genes **(A)** and number of significant associations to phenotypes **(B)**.

**(C)** Average number of significant associations to phenotypes between clusters enriched for gene ontology and clusters not enriched for gene ontology (Wilcoxon Rank Test).

**(D-F)** Conservation of clusters across permutations in normalisation method. Degree of conservation between clusters identified using CLR normalisation and **(D)** no normalisation, **(E)** relative count normalisation, and **(F)** log normalisation. (Pearson's correlation).

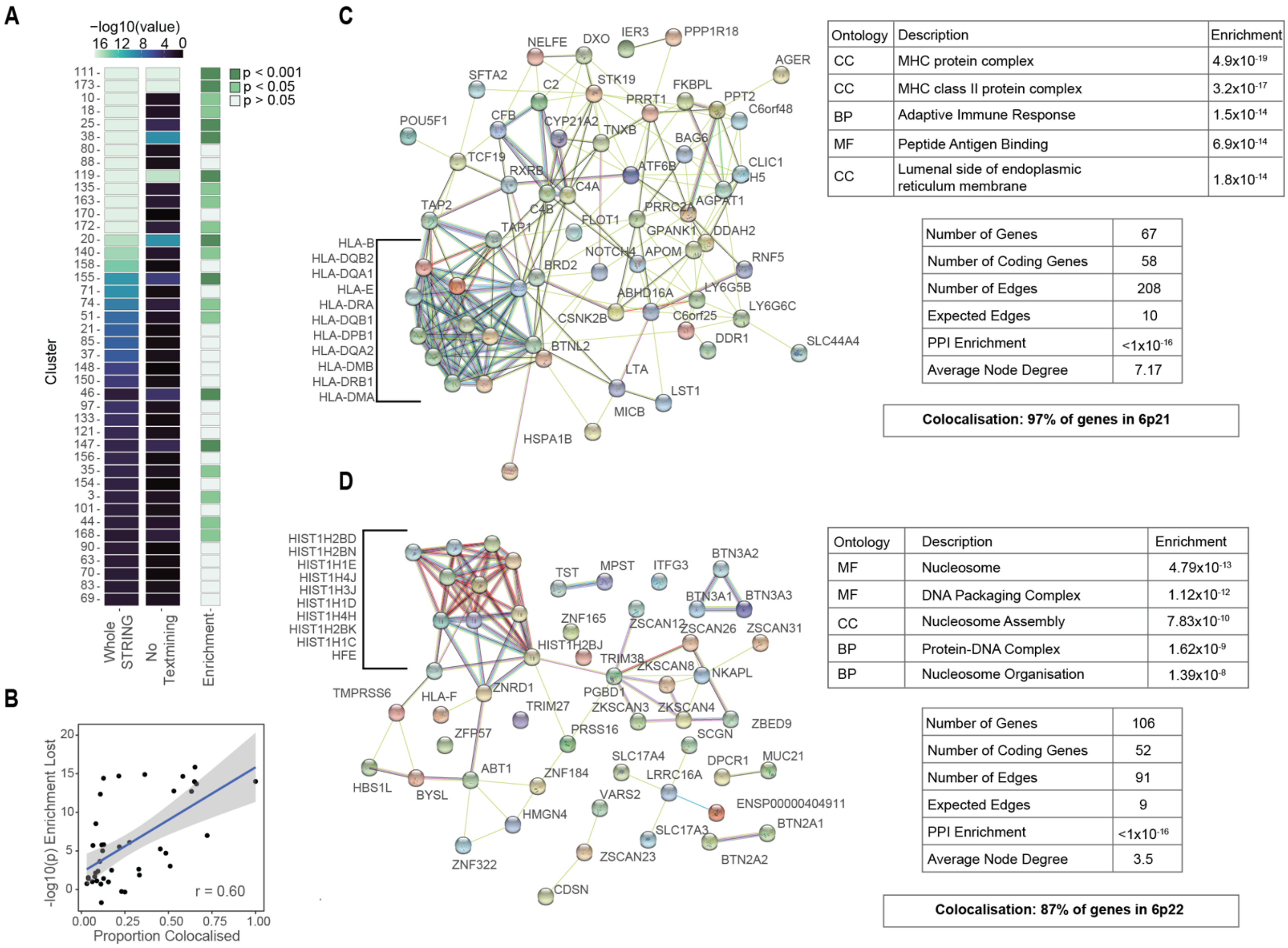


**Supplementary Figure 6. Genes in highly colocalised clusters have related biological functions.**

**(A)** Significance of STRING protein-protein interaction enrichment of clusters using all components at a medium confidence threshold and without the text-mining component at a low confidence threshold. The enrichment column indicates whether the cluster remains enriched after removal of the text mining component.

**(B)** Relationship between the amount of enrichment lost and the degree of colocalization in a cluster. Modelled using a linear model (Pearson's correlation = 0.60).

**(C, D)** Breakdown of STRING protein-protein interaction, gene ontology enrichment and colocalization of **(C)** cluster 173 and **(D)** cluster 111.

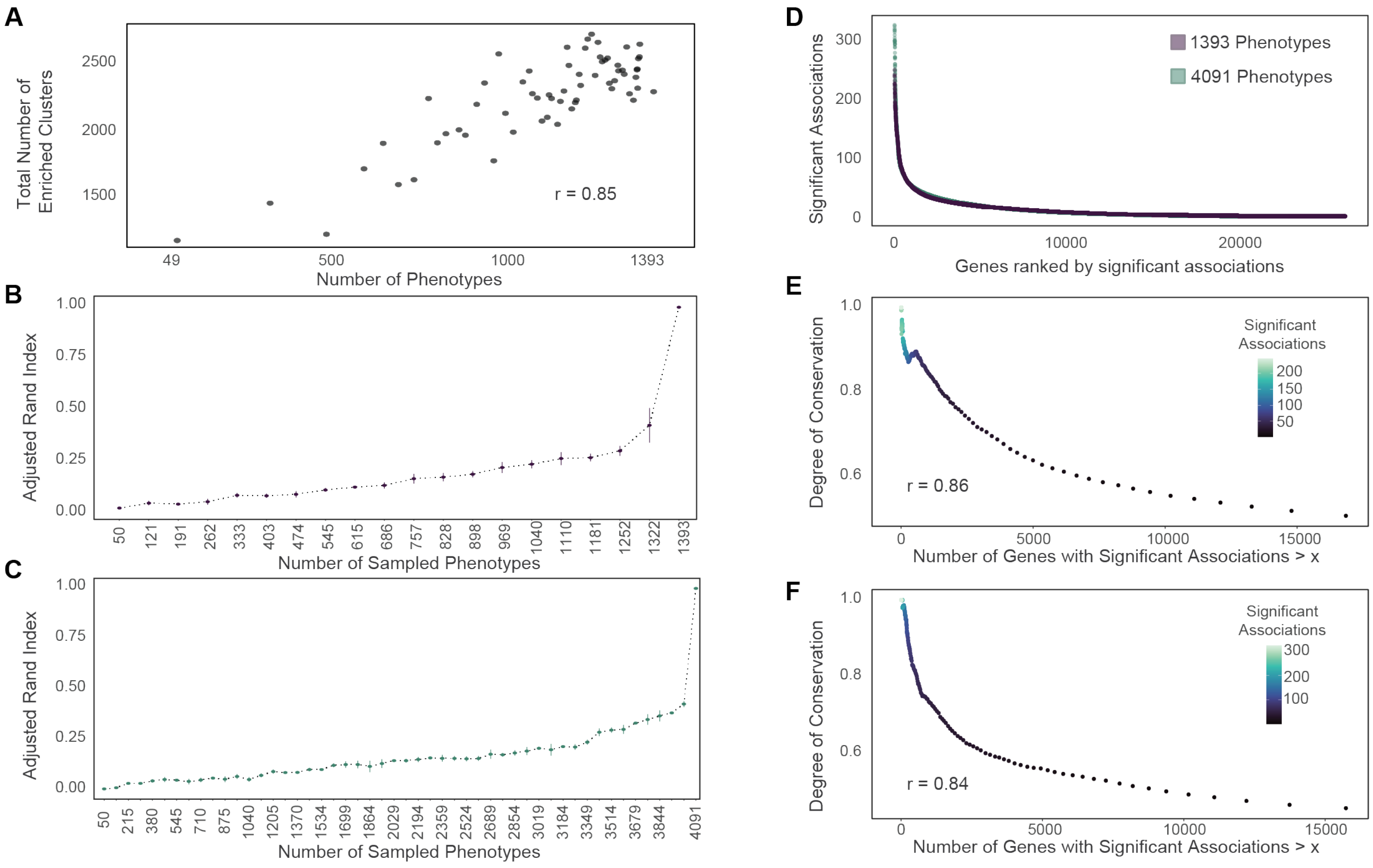


**Supplementary Figure 7. Phenotype subsampling reveals data is unsaturated**

**(A)** Effect of decreasing phenotype number on the total number of enriched clusters, Pearson's correlation was used to determine the correlation.

**(B-C)** Phenotypes were subsampled 5 times for each of **(B)** 20 equal intervals between 50 and 1393 and **(C)** 50 equal intervals between 50 and 4091. The UnTANGLeD clustering pipeline was applied to each supplied dataset. Adjusted rand index was calculated for each sub-sampled dataset compared with clustering computed using **(B)** 1393 phenotypes **(C)** 4091 phenotypes.

**(D)** The number of significant associations identified by MultiXcan for each gene from a dataset containing 1393 phenotypes and a dataset containing 4091 phenotypes.

**(E-F)** Relationship between the number of significant associations a gene has and its degree of conservation between the maximum number of phenotypes and the first replicate of the subsampling trial with **(E)** 1322 **(F)** 4009 phenotypes. Genes were grouped into bins by selecting genes which possessed more significant associations than x, where x ranged from **(F)** 1 to 246 in increments of 1 **(F)** 1 to 322 in increments of 1. Bins are represented by the number of genes remaining with more significant associations than x.

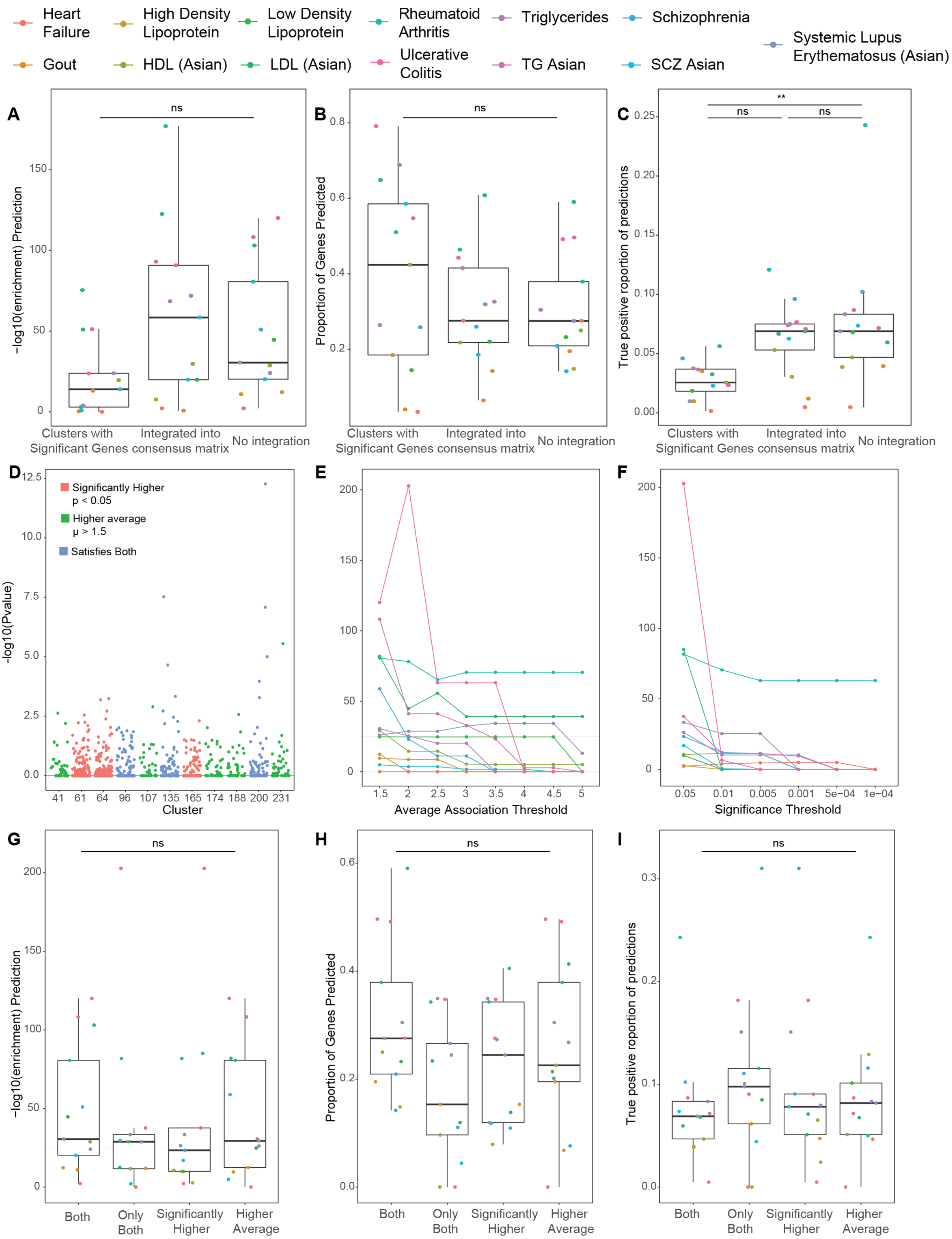
Heart Failure
High Density Lipoprotein
Low Density Lipoprotein
Rheumatoid Arthritis
Triglycerides
Schizophrenia
Systemic Lupus Erythematosus (Asian)
Gout
HDL (Asian)
LDL (Asian)
Ulcerative Colitis
TG Asian
SCZ Asian
A
ns
−log10(enrichment) Prediction
0
50
100
150
Clusters with Significant Genes
Integrated into consensus matrix
No integration
B
ns
Proportion of Genes Predicted
0.2
0.4
0.6
0.8
Clusters with Significant Genes
Integrated into consensus matrix
No integration
C
**
ns
ns
True positive roportion of predictions
0.00
0.05
0.10
0.15
0.20
0.25
Clusters with Significant Genes
Integrated into consensus matrix
No integration
D
Significantly Higher p < 0.05
Higher average μ > 1.5
Satisfies Both
-log10(Pvalue)
0.0
2.5
5.0
7.5
10.0
12.5
41 61 64 96 107 135 165 174 188 200 231
Cluster
E
0
50
100
150
200
1.5 2 2.5 3 3.5 4 4.5 5
Average Association Threshold
F
0
50
100
150
200
0.05 0.01 0.005 0.001 5e−04 1e−04
Significance Threshold
G
ns
−log10(enrichment) Prediction
0
50
100
150
200
Both
Only Both
Significantly Higher
Higher Average
H
ns
Proportion of Genes Predicted
0.0
0.2
0.4
0.6
Both
Only Both
Significantly Higher
Higher Average
I
ns
True positive roportion of predictions
0.0
0.1
0.2
0.3
Both
Only Both
Significantly Higher
Higher Average

**Supplementary Figure 8. Testing of integration methods and selection parameters for predicting which clusters are associated with a trait.**

**(A-C)** (*$P < 0.05$) Comparison of selecting clusters that contain significant genes from the trait, integrating the phenotype into the clustering pipeline and using the complete association signature on prediction enrichment **(A),** proportion of associated genes predicted **(B),** and proportion of predictions that were a true positive **(C)**.

**(D)** Representative image of selection criteria based on either significance or average association.

(**E-F)** Effect of different significance **(E)** and average association **(F)** thresholds for implicating a cluster in an independent phenotype on the prediction enrichment.

**(G-I)** Effect of using only the significance threshold, only the average association, or both as a selection criterion on the prediction enrichment **(G)**, proportion of associated genes predicted **(H)** and proportion of predictions that were a true positive **(I)**. We used a Kruskal Wallis one way analysis of variance, followed by a Dunn's Test to determine any significant differences.

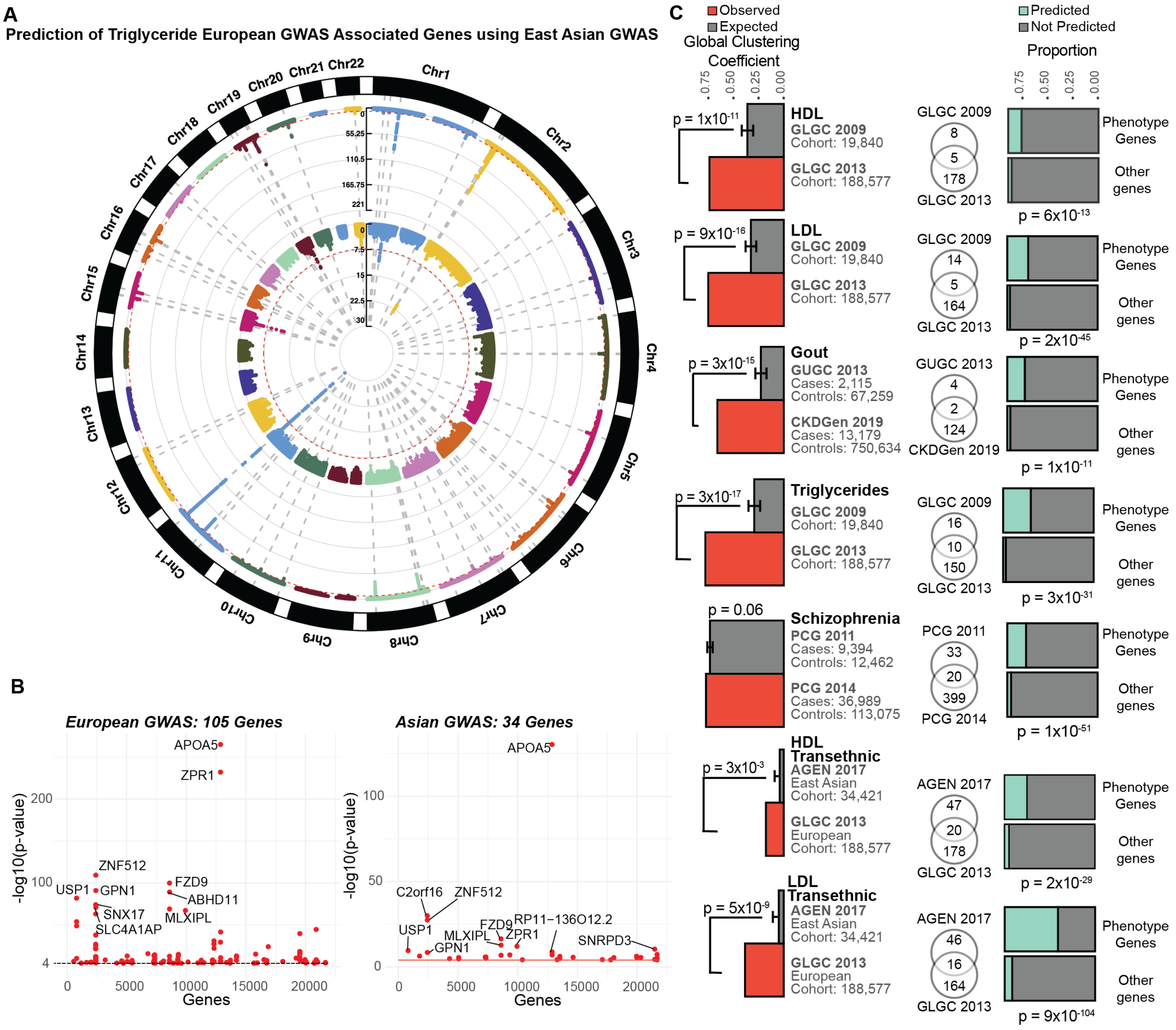


**Supplementary Figure 9. Prediction of novel GWAS genes using a GWAS with a smaller sample size.**

**(A)** Manhattan plot of loci identified by a European and East Asian GWAS of Triglyceride levels.

**(B)** Manhattan plot of S-MultiXcan genes for the European and East Asian GWAS of triglyceride levels respectively, genes are ordered according to their genomic positions.

**(C)** Increase in global clustering coefficient and prediction enrichment across 4 European and 3 Transethnic phenotype pairs as calculated using bootstrapping and chi-squared enrichment test.

**Supplemental Table 1.** Summary of independent studies used to perform gene prediction and prioritization analyses. Phenotype, source study, cohort sizes, cohort ancestry, significant loci and genes provided.

| Trait | Study | Year | Consortium | Cohort Size | Control | Case | Loci | MultiXcan Genes | Ethnicity |
|---|---|---|---|---|---|---|---|---|---|
| Gout | Köttgen *et al* | 2013 | GUGC | 69,374 | 2,115 | 67,259 | 3 | 4 | European |
| | Tin *et al* | 2019 | CKDGen | 763,813 | 13,179 | 750,634 | 32 | 124 | European & African American |
| Heart Failure | Sudlow *et al* | 2015 | UK Biobank | 361194 | 359,789 | 1,405 | 5 | 5 | European |
| | Shah *et al* | 2020 | HERMES | 977,323 | 930,014 | 47,309 | 7 | 29 | European |
| High Density Lipoprotein | Kathiresan *et al* | 2009 | GLGC | 19,840 | - | - | 3 | 8 | European |
| | Willer *et al* | 2013 | GLGC | 188,577 | - | - | 66 | 178 | European |
| | Spracklen *et al* | 2017 | AGEN | 34,421 | - | - | 9 | 47 | East Asian |
| Low Density Lipoprotein | Kathiresan *et al* | 2009 | GLGC | 19,840 | - | - | 4 | 14 | European |
| | Willer *et al* | 2013 | GLGC | 188,577 | - | - | 50 | 164 | European |
| | Spracklen *et al* | 2017 | AGEN | 34,421 | - | - | 7 | 46 | East Asian |
| Rheumatoid Arthritis | Stahl *et al* | 2010 | - | 25,708 | 20,169 | 5,539 | 10 | 110 | European |
| | Okada *et al* | 2014 | - | 79,799 | 60,565 | 19,234 | 45 | 296 | European & East Asian |
| Schizophrenia | Schizophrenia Psychiatric Genome-Wide Association Study (GWAS) Consortium | 2011 | PCG | 21,856 | 9,394 | 12,462 | 11 | 33 | European |
| | Schizophrenia Working Group of the Psychiatric Genomics Consortium | 2014 | PCG | 150,064 | 36,989 | 113,075 | 100 | 399 | European |
| | Lam *et al* | 2019 | PCG | 58,140 | 22,778 | 35,362 | 12 | 152 | East Asian |
| Systemic Lupus Erythematosus | Bentham *et al* | 2015 | | 14,267 | 5,201 | 9,066 | 38 | 176 | European |
| | Wang *et al* | 2021 | | 12,653 | 4,222 | 8,431 | 30 | 178 | Han Chinese |
| Triglycerides | Kathiresan *et al* | 2009 | GLGC | 19,840 | - | - | 3 | 16 | European |
| | Willer *et al* | 2013 | GLGC | 188,577 | - | - | 39 | 150 | European |
| Ulcerative Colitis | Spracklen *et al* | 2017 | AGEN | 34,421 | - | - | 8 | 34 | East Asian |
| | Anderson *et al* | 2011 | - | 26,405 | 19,718 | 6,687 | 29 | 153 | European |
| | de Lange et al | 2017 | - | 45,975 | 33,609 | 12,366 | 80 | 278 | European |

**Supplemental Data in Supplementary Tables**

**Data S2. Supplementary Table 2**
Details of all phenotypes used to construct the gene-trait association matrix. Cohort size, study cohort, phenotype category and number of significant S-MultiXcan associations provided.

**Data S3. Supplementary Table 3**
Summary details of the 173 UnTANGLeD clusters for average silhouette scores, correlation scores, enrichment profiles across gene ontology, disease ontology, KEGG pathways, Tissue specificity (TissueEnrich) and colocalization within a gene segment. Full gene list for each cluster provided.

**Data S4. Supplementary Table 4**
All gene ontology enriched terms for each cluster provided. Enrichment performed by *ClusterProfiler*, FDR corrected p-value < 0.01.

**Data S5. Supplementary Table 5**
All KEGG pathway enriched terms for each cluster provided. Enrichment performed by *ClusterProfiler*, FDR corrected p-value < 0.01.

**Data S6. Supplementary Table 6**
All disease ontology enriched terms for each cluster provided. Enrichment performed by *ClusterProfiler*, FDR corrected p-value < 0.01.

**Data S7. Supplementary Table 7**
Top tissue with greatest fold enrichment for each cluster provided. Enrichment performed by *TissueEnrich.*

**Data S8. Supplementary Table 8**
All WikiPathways enriched terms for each cluster provided. Enrichment performed by *ClusterProfiler*, FDR corrected p-value < 0.01.

**Data S9. Supplementary Table 9**
The top 5 enriched terms for each UnTANGLeD cluster from each EnrichR library. Enrichment performed using *EnrichR*. Multiple testing corrected p-value < 0.01.

**Data S10. Supplementary Table 10**
Protein families and domains from Pfam Domain (2019) Database with more than one member in a single cluster. Enrichment calculated using *EnrichR*.

**Data S11. Supplementary Table 11**
Top enriched genomic segment for each cluster provided. Enrichment performed by *ClusterProfiler*, FDR corrected p-value < 0.01.
